# Potential evolutionary body size reduction in a Malagasy primate (*Propithecus verreauxi*) in response to human size-selective hunting pressure

**DOI:** 10.1101/2020.03.23.004234

**Authors:** Alexis P. Sullivan, Laurie R. Godfrey, Richard R. Lawler, Heritiana Randrianatoandro, Laurie Eccles, Brendan Culleton, Timothy M. Ryan, George H. Perry

## Abstract

The Holocene arrival of humans on Madagascar precipitated major changes to the island’s biodiversity. The now-extinct, endemic “subfossil” megafauna of Madagascar were likely hunted by the island’s early human inhabitants. Perhaps in part due to preferential hunting of larger prey, no surviving species on Madagascar is larger than 10 kg. Outside of Madagascar, size-selective hunting pressure has resulted in the phyletic dwarfism of many still-living species across a diversity of phyla. On Madagascar, some subfossil bones of extant lemurs are considerably larger than those of the modern members of their species, but relatively large distances between the subfossil localities and modern samples that have been compared to date makes it impossible to reject the possibility that these size differences more simply reflect pre-existing ecogeographic variation. Here, we used high-resolution 3D scan data to conduct comparative morphological analyses of subfossil and modern skeletal remains of one of the larger extant lemurs, Verreaux’s sifakas (*Propithecus verreauxi*) from subfossil and modern sites ∼10 km adjacent: Taolambiby (bones dated to 725-560 – 1075-955 cal. years before present) and Beza Mahafaly Special Reserve, respectively. We found that the average subfossil sifaka bone (n=12) is 9% and significantly larger than that of modern sifakas (n=31 individuals; permutation test; p=0.037). When restricting the analysis to the single element and side with the largest representation in the subfossil sample (n=4 right distal femora), the average subfossil bone is 10% larger (p=0.046). While we cannot yet conclude whether this size difference reflects evolutionary change or an archaeological aggregation/taphonomic process, if this is a case of phyletic dwarfism in response to human size-selective harvesting pressures then the estimated rate of evolutionary change is slightly higher than that previously calculated for other archaeological cases of this phenomenon.

## INTRODUCTION

Intensive human behaviors including harvesting and predation, landscape modification, and translocation have effected non-human morphological evolution for at least tens of thousands of years [1]. For example, size-selective hunting pressure by humans has resulted in documented body size or feature reduction for many different non-human taxa [2,3], from aquatic invertebrates and vertebrates such as snails [4,5] and salmon [6,7] to terrestrial mammals like bighorn sheep [8,9]. For terrestrial taxa, assessments of body size diminution due to human hunting pressures have been largely restricted to ungulates [2,3]; no such process has yet been recorded for a non-human primate species.

The Malagasy megafauna were comprised of at least 28 large-bodied species from ∼11 kg to ∼650 kg in size [10], including at least 17 lemurs (primates), the largest of which had an estimated body mass of ∼160 kg [11,12]. The timing of human arrival and permanent residence on Madagascar is uncertain [13,14] but may extend to ∼10,500 years BP or earlier [15–17]. There are reports of intentional human processing marks on lemur bones in southwest Madagascar dating to ∼2,300 years BP [11,18,19]. Human harvesting pressures, perhaps including preferential hunting of larger animals, have often been discussed as potential contributing factors in population declines and eventual extinctions of the island’s megafauna [18,20–23]. Now, no surviving endemic terrestrial vertebrate species on Madagascar has a body mass larger than 10 kg [24].

On Madagascar, subfossil bones of still-living lemur species recovered from archaeological and paleontological contexts are often considerably larger than those of their modern counterparts [19,25– 27]. Godfrey et al. [26] noted that there are many subfossil sites in the southwest with *Propithecus verreauxi* bones that are both more robust and longer than modern *Propithecus* in the same region. Indeed Lamberton [28] had assigned a new species nomen, *Propithecus verreauxoides*, to subfossil specimens from Tsirave (south central Madagascar) because of its larger sizes for major skull measurements and long bone lengths relative to those of modern *Propithecus verreauxi*. Muldoon and Simons [25] reported a similar size disparity between *Lepilemur* subfossils from Ankilitelo and extant *Lepilemur* from the same general region, but they also noted that prehistoric-extant size differences could simply reflect ecogeographic variation rather than recent body size evolution.

For our present study, we identified an opportunity to investigate a potential case of non-human primate phyletic dwarfism due to human hunting pressures without the complicating factor of large geographic distance between where the compared subfossil and modern individuals lived. Specifically, we report the results from a morphological analysis of Verreaux’s sifaka (*Propithecus verreauxi*) postcranial skeletal remains from the subfossil Taolambiby site and the Beza Mahafaly Special Reserve, located ∼10 km apart (**Figure 1**), to test the hypothesis that *P. verreauxi* body size decreased following the earliest appearance of cut-marked bones.

**Figure 1:**
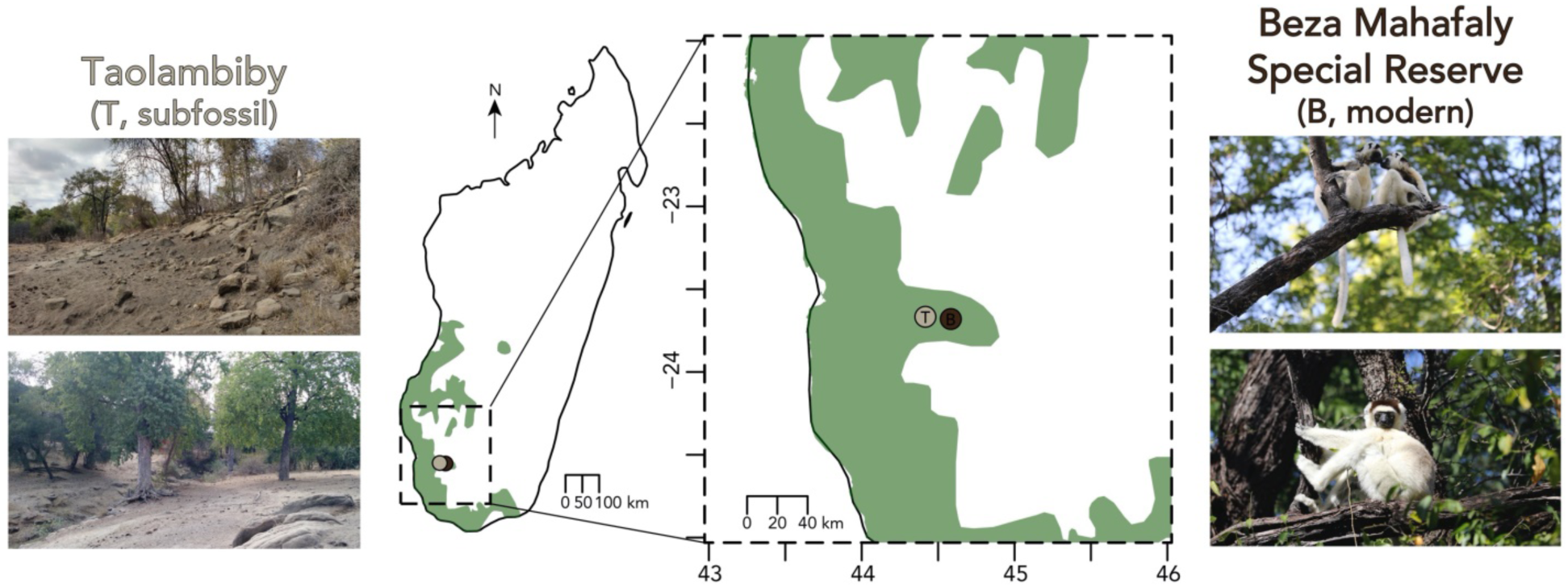
*Propithecus verreauxi* osteological collection sites. Verreaux’s sifaka range data in green from IUCN Red List database [species 18354], last updated 2014. Subfossil osteological materials were collected from Taolambiby (tan dot), and modern remains from the Beza Mahafaly Special Reserve (brown dot).

## MATERIALS AND METHODS

### Osteological Collections

We surveyed the osteological collections from two sites in southwestern Madagascar in an attempt to characterize a body size diminution in local *Propithecus verreauxi* over time. *P. verreauxi* has an adult body mass ranging from ∼2.5 – 3.5 kg [29,30], and has historically been hunted by humans across multiple parts of its range [31–33]. Specifically, more than 250 *P. verreauxi* skeletal and craniodental elements were surface-collected from along the Taolambiby village river wash by Alan Walker in 1966 [17,19] (**Figure 1**). This Taolambiby collection was donated to the Anthropological Primate Collection at the University of Massachusetts, Amherst (UM-TAO). Cut marks, chop marks, and/or spiral fractures indicative of human processing were identified on 62% of the *P. verreauxi* elements [19]. The Taolambiby subfossil material is fragmentary; from the larger collection available to us there were a total of 15 elements that could be identified confidently as adult *P. verreauxi* femora and humeri that we included in our study.

Immediately adjacent to this subfossil site/collection is the Beza Mahafaly Special Reserve (BMSR; **Figure 1**), which is home to an extant sifaka population that has been monitored and studied since 1984 [34]. In the first 25 years of BMSR collection and management of sifaka long-term data, 718 individuals have been captured, measured, and marked; now there are >900 individuals [34]. Since 1985, researchers have also been collecting and labeling faunal skeletal remains discovered in the course of observation or survey [35]. The resulting Beza Mahafaly Osteological Collection (BMOC) is comprised of skeletal elements from all four extent lemur species living in the area, along with invasive wildcats. *P. verreauxi* was represented in this collection by the remains of 31 adult individuals at the time of our data collection (see **Supplementary Table 1** for full list of individuals).

### Data Collection

We focused on femora and humeri because there is a strong positive correlation between measurements from these long bones and overall body size in *Propithecus* lemurs [36]. Juvenile specimens, identified by the presence of an epiphyseal line or an unfused epiphysis [37], were excluded from our analysis. Of the modern BMOC adults (n=31 total), sex was known for eight individuals (26%; n=4 males, n=4 females) based on the association of their remains with collars and/or other identifying features (see **Supplementary Table 1**). Since sifakas are not sexually dimorphic [29,38– 40], male and female adults are expected to have approximately similar body sizes and sex was not included as a variable in our analyses.

We collected 3D surface data from every adult *P. verreauxi* long bone that was available in each collection, fragmented or whole, right or left, with a portable Artec Space Spider (Artec 3D, Luxembourg; **Figure 2**; **Table 1**). All data collection at the Beza Mahafaly Special Reserve was approved by the Madagascar National Parks organization. The Spider scanner records high-resolution geometry and texture data at up to 0.1 mm resolution and 3D point accuracy up to 0.5 mm. Each of the 106 adult skeletal elements from these collections (n=94 modern, 15 subfossil) was affixed to a turntable, and the Artec Space Spider was used to collect between 4-8 scans from multiple angles to capture the entirety of the bone. Linear caliper measurements (n=5 for humeri, 6 for femora; **Table 1**) were also collected from every bone (see **Supplementary Table 2**).

**Table 1:** Morphometric caliper and 3D surface scan measurement descriptions.

|  | Measurement | Measurement Description | Number of Elements Measured |  |  |  |  |  |  |  |
| --- | --- | --- | --- | --- | --- | --- | --- | --- | --- | --- |
|  |  |  | 3D Scanner |  |  |  | Radial Caliper |  |  |  |
|  |  |  | Modern<br>Right | Modern<br>Left | Subfossil<br>Right | Subfossil<br>Left | Modern<br>Right | Modern<br>Left | Subfossil<br>Right | Subfossil<br>Left |
| **Humerus** | MHL | Maximum length measured between top of humeral head and most distant point on distal humerus | 18 | 22 | 0 | 0 | 16 | 19 | 0 | 0 |
|  | MHD1 | Maximum midshaft humeral diameter, measured just below deltoid | 23 | 23 | 0 | 0 | 23 | 20 | 0 | 0 |
|  | MHD2<br>[3D only] | Minimum midshaft humeral diameter, measured just below deltoid | 23 | 23 | 0 | 0 | NA | NA | NA | NA |
|  | VHD | Vertical head diameter, superoinferior diameter | 21 | 23 | 1 | 0 | 20 | 20 | 1 | 0 |
|  | HHW | Humeral head width, external transverse mediolateral diameter | 21 | 23 | 1 | 0 | 20 | 20 | 1 | 0 |
|  | BBH | Biepicondylar breadth, greatest distance between medial and lateral epicondyles, parallel to humeral shaft | 22 | 23 | 0 | 0 | 21 | 20 | 0 | 0 |
|  | HHSa [3D only] | Avizo-calculated humeral head surface area | 21 | 23 | 1 | 0 | NA | NA | NA | NA |
| **Femur** | MFL | Maximum length that can be measured between the top of the greater trochanter and bottom of the most distal condyle | 20 | 19 | 0 | 0 | 21 | 18 | 0 | 0 |
|  | MFD | Anteroposterior (sagittal) midshaft diameter | 23 | 24 | 0 | 0 | 24 | 24 | 0 | 0 |
|  | FMD<br>[3D only] | Mediolateral (transverse) midshaft diameter | 23 | 24 | 0 | 0 | NA | NA | NA | NA |
|  | MFC<br>[caliper only] | Midshaft circumference | NA | NA | NA | NA | 23 | 24 | 0 | 0 |
|  | FHH | Femoral head height, superoinferior diameter | 24 | 20 | 2 | 2 | 24 | 20 | 2 | 2 |
|  | FHW | Femoral head width, anteroposterior diameter | 24 | 20 | 2 | 2 | 24 | 20 | 2 | 2 |
|  | FBE | Femoral biepicondylar breadth, distance between medial-most and lateral-most points on epicondyles | 23 | 21 | 2 | 1 | 23 | 21 | 1 | 2 |
|  | DML1<br>[3D only] | Width of medial distal condyle | 23 | 21 | 0 | 2 | NA | NA | NA | NA |
|  | DML2<br>[3D only] | Width of lateral distal condyle | 23 | 21 | 4 | 3 | NA | NA | NA | NA |
|  | FHSA<br>[3D only] | Avizo-calculated femoral head surface area | 24 | 20 | 2 | 2 | NA | NA | NA | NA |
|  | DSA<br>[3D only] | Avizo-calculated condylar surface area | 23 | 21 | 0 | 1 | NA | NA | NA | NA |

Note: Unless specified, each measurement was collected with both linear calipers and 3D scan data.

**Figure 2:**
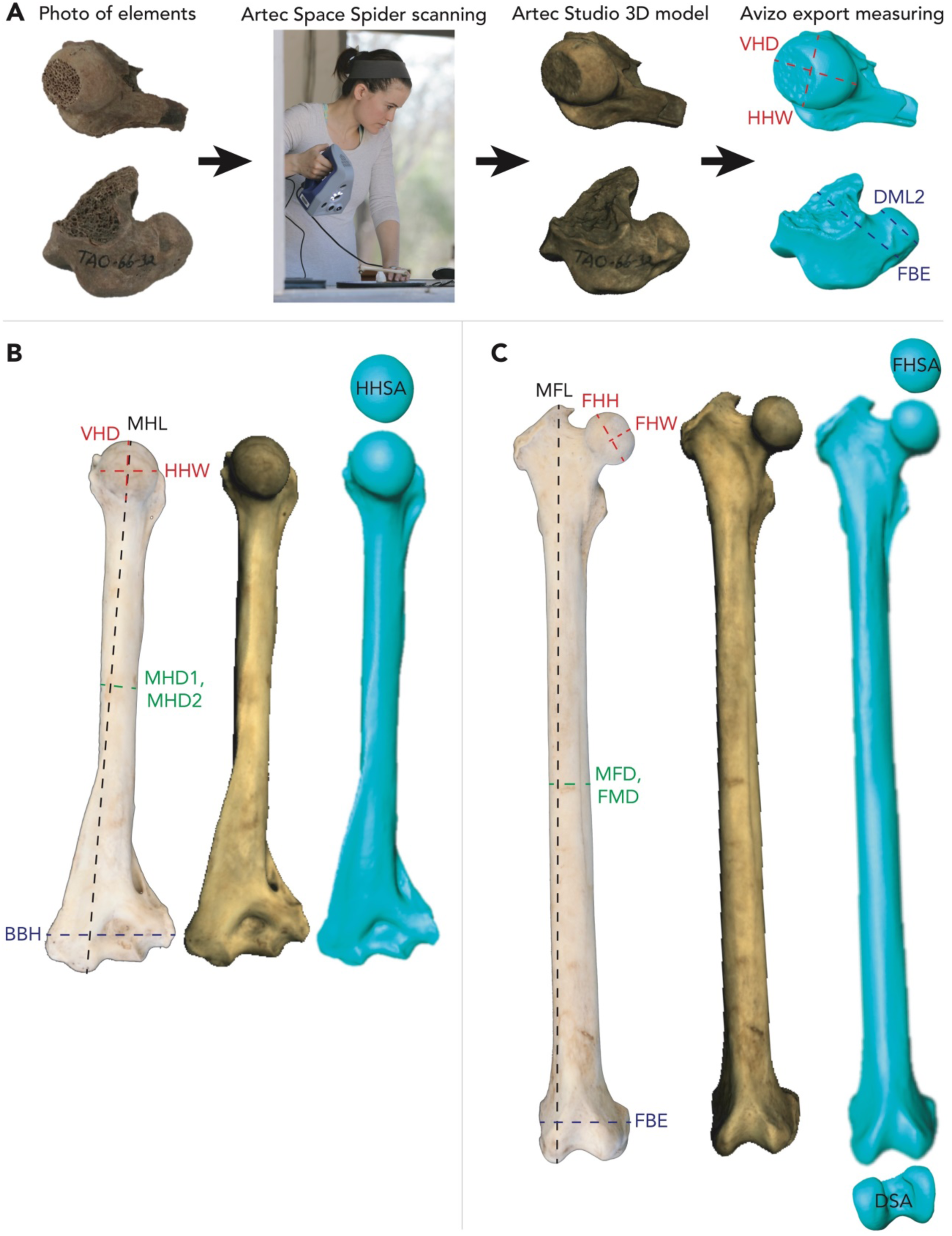
Examples of osteological elements of *P. verreauxi* and the measurements collected for this study. Descriptions of each measurement are available in **Table 1**. (A) Two subfossil elements from Taolambiby. The humeral head (UM-TAO-66-2) was radiocarbon dated to ∼1000 BP and the distal femur (UM-TAO-66-32) was dated to ∼740 BP. Diagram of measurements collected from (B) humeri and (C) femora (distal condyle widths not pictured; example humerus and femur from BMOC-020).

### 3D Surface Scan Post-Processing

The surface scan data were post-processed with Artec Studio 11 software to form 3D models for each element. Specifically, scans were first individually cleaned with the “Eraser” function to remove the turntable and other background noise. The multiple scans of the same element were then aligned to one another with at least three points of common geometry. The aligned scan data were then registered (“Global Registration” with settings for 50 mm minimal distance and 5000 iterations) and fused (“Fusion”: default “Outlier Removal” and “Sharp Fusion” settings with “Watertight” 0.3 mm resolution). 3D meshes of each element were then exported and individually measured as indicated in **Table 1** and **Figure 2** with Avizo 9.4 software (see **Supplementary Table 3**). We measured maximum length, midshaft diameters and circumferences for all whole femora, as well as femoral head heights, widths, and surface areas, bi-epicondylar breadth, and condylar widths and surface areas when available [36,37,41]. We measured maximum length and midshaft diameters for all whole humeri, as well as humeral head diameters, widths, and surface areas, and bi-epicondylar breadth when available [36,37,41]. Surface areas were determined by isolating the osteological region of interest from the rest of the bone, then using the Materials Surface Area Statistics tool available in Avizo. All 3D models are available on MorphoSource (“Sullivan/Perry Lab *Propithecus verreauxi* Surface Scans” Project <u>ID 698</u>).

### AMS Radiocarbon Dating

It was important to establish the antiquity of each subfossil element included in our analysis with radiocarbon ^14^C dating methods. Of 18 Taolambiby *P. verreauxi* skeletal elements dated in a prior study, one (5.6%) was modern and the remaining 17 (94.4%) ranged from 605 to 1185 cal BP [42, Supplementary Table 3]. Of those 17 previously dated, non-modern specimens, one was a proximal humerus and seven were femora. The humeral fragment and five of the femoral fragments were sufficiently intact for us to obtain at least one size measurement per bone (**Table 2**).

**Table 2:** Subfossil *P. verreauxi* elements that were included in this study and their corresponding radiocarbon dates.

| Sample ID | Element (Side, End) | Human Processing Mark(s) | <sup>14</sup>C Lab ID | Conventional <sup>14</sup>C Age (BP) | Calibrated 95.4% CI (cal. BP) | Reference |
| --- | --- | --- | --- | --- | --- | --- |
| UM-TAO-66-26 | Femur (R, D) | Chop | PSUAMS-2480 | 1010 ± 15 | 925 - 800 | This study |
| UM-TAO-66-29 | Femur (R, D) | Chop | PSUAMS-2479 | 865 ± 20 | 770 - 680 | This study |
| UM-TAO-66-32 | Femur (R, D) | Chop | N/A* | 740 ± 40 | 725 - 560 | This study |
| UM-TAO-66-33 | Femur (R, D) | Chop | PSUAMS-2477 | 975 ± 20 | 920 - 790 | This study |
| UM-TAO-66-31 | Femur (L, D) | Chop | N/A* | 850 ± 35 | 775 - 670 | This study |
| UM-TAO-66-34 | Femur (L, D) | Chop, slice through condyles | PSUAMS-2476 | 840 ± 20 | 740 - 675 | This study |
| UM-TAO-66-2 | Humerus (R, P) | Chop, cut marks | CAMS-147111 | 1035 ± 25 | 960 - 800 | [42] |
| UM-TAO-66-21 | Femur (R, P) | Chop | CAMS-147036 | 1130 ± 30 | 1060 - 930 | [42] |
| UM-TAO-66-23 | Femur (L, P) | Chop | CAMS-147628 | 1055 ± 30 | 965 - 805 | [42] |
| UM-TAO-66-24 | Femur (L, P) | Chop | CAMS-147114 | 940 ± 40 | 915 - 730 | [42] |
| UM-TAO-66-25 | Femur (R, P) | Chop | CAMS-147113 | 915 ± 35 | 905 - 685 | [42] |
| UM-TAO-66-30 | Femur (L, D) | Chop | CAMS-147332 | 1165 ± 30 | 1075 - 955 | [42] |

\*UM-TAO-66-31 and -32 were not assigned PSUAMS-# because there is no EA-IRMS data (-31) or poor C:N ratio (-32).

Nine additional (not previously radiocarbon dated) adult *P. verreauxi* femora from Taolambiby were also available for possible inclusion in our analysis. For each of these specimens, we first collected all measurements and 3D surface scan data before sampling 200-500 mg from the subfossil skeletal remains for AMS radiocarbon ^14^C dating and stable isotope analyses at the Penn State University Human Paleoecology and Isotope Geochemistry Laboratory. The bone samples were scraped with blades to remove adhering material and clipped into small pieces. As a precaution, we removed possible conservants and adhesives by sonicating the scraped bone samples in washes of ACS-grade methanol, acetone, and dichloromethane for 20 minutes each at room temperature. Bone collagen was extracted and purified after sonication. Samples were demineralized for 1-3 days in 0.5N hydrochloric acid at 5°C. The demineralized pseudomorph was rinsed twice in 18.2Ω/cm Nanopure water for 20 minutes. The pseudomorph was gelatinized for 10 hours at 60°C in 0.01 Normal hydrochloric acid. The resulting gel was then lyophilized and weighed to determine percent yield as a first evaluation of the degree of bone collagen preservation.

In this case the collagen samples were relatively poorly preserved and so they were pretreated using a modified XAD process [43,44] after demineralization and gelatinization. The gelatin was hydrolyzed in 1.5 mL of 6 Normal hydrochloric acid for 24 hours at 110°C. Supelco ENVI-Chrome P SPE (Solid Phase Extraction) columns were fitted with a Millex HV PVDF 0.45 µm filter unit, and both were equilibrated with 50 mL of 6 Normal hydrochloric acid. The 1.5 mL sample hydrolyzate was pipetted into the SPE column and driven through with a syringe and an additional 10 mL of 6 N hydrochloric acid dropwise into a prepared 20 mm culture tube. The hydrolyzate (now bone collagen amino acids) was dried into a viscous syrup by passing UHP nitrogen gas over the heated (50°C) sample for about 8 hours.

The XAD amino acids were analyzed for carbon and nitrogen concentrations and stable isotope ratios at the Yale Analytical and Stable Isotope Center with a Costech elemental analyzer (ECS 4010) and Thermo DeltaPlus isotope ratio mass spectrometer. Sample quality was evaluated by %C, %N, and the C:N ratio before AMS ^14^C dating. Good quality amino acid samples were then weighed (3.5-4.5 mg) into 8” quartz tubes, with 60 mg CuO and a ∼2mm snip of 1mm diameter 99.9% silver wire, then sealed under vacuum and combusted at 800°C for 3 hours.

The resulting CO_2_ was reduced to graphite at 550°C using UHP hydrogen gas and an iron catalyst, with the reaction water removed by magnesium perchlorate (Mg(ClO_4_)_2_). Graphite samples were pressed into targets and loaded onto a target wheel with oxalic acid (OXII) primary standards, known age bone secondaries and ^14^C free Pleistocene whale blank, and measured on a modified NEC 1.5SDH-1 500kV compact accelerator mass spectrometer housed in the Penn State Earth and Environmental Sustainability Laboratories. Three subfossil elements, UM-TAO-66-22, UM-TAO-66-28, and an unaccessioned femoral head, had insufficient remaining collagen for either isotopic or radiocarbon analysis, and thus were not included in our subsequent morphological analyses. Two samples were run on the AMS but not assigned PSUAMS lab codes, and should be considered provisional. C and N abundance and isotope ratios were unavailable for UM-TAO-66-31, and C:N ratio for UM-TAO-66-32 was 3.64, at the limits of the acceptable range. AMS radiocarbon results from this study and previous dates by Crowley and Godrey [42] were calibrated with OxCal v. 4.3.2 [45] using the SHCal13 southern hemisphere curve [46] and are presented in **Table 2**.

### Correlations Between 3D Scan and Caliper Measurements

We directly compared the equivalent raw caliper and 3D scan measurements to each other as an accuracy check for each measurement technique. For every available measurement category (MFL, MFD, FHH, FHW, FBE; MHL, MHD1, VHD, HHW, BBH) we calculated the percent difference between the corresponding caliper measurement and 3D scan measurement taken for each individual skeletal element:

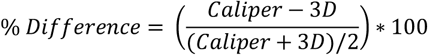

We determined that, on average, the caliper measurements were 0.68% smaller than the equivalent 3D measurements for the modern skeletal remains and 1.20% larger for the subfossil elements (**Supplemental Table 4;** see **Supplemental Table 5** for all caliper versus 3D measurement comparisons). This difference is likely due to difficulties in identifying measurement landmarks with the calipers on the more broken subfossil materials.

With the advantages of increased maximum/minimum measurement accuracy and surface area calculation tools, more 3D model measurements were able to be collected from the fragmented modern and subfossil materials relative to those with the caliper: 10 femoral 3D measurements versus 6 with caliper, and 7 humeral 3D versus 5 with caliper. For some bones, 3D digital measurement only was possible. Therefore, all results presented here are based on the 3D model measurements to increase the number of specimens included in the analyses.

### 3D Surface Scan Measurement Analyses

The limited number of 3D surface scan measurements that could be taken on each subfossil bone precluded our ability to directly estimate individual body sizes. Therefore, we compared the individual bone measurements to each other. To do so, for each measurement we first calculated the geometric mean (geoMean, *GM*; [47]) from all available modern individuals for that measurement (**Table 3, Step A**). As an example, the geoMean of the modern right femoral head height (FHH) from the 3D surface scan data was 12.37 mm. These geoMeans were then used to calculate a relative fold change (*FC*) value for each individual modern and subfossil skeletal element (**Table 3, Step B**). For example, the FHH measurements for modern BMOC-001 was 12.21 mm and for subfossil UM-TAO-66-25 was 12.90 mm. The FC for each of these individuals was calculated from the 12.37 mm GM: -0.01 and 0.04, respectively.

**Table 3:** Measurement analysis calculations and examples.

| Step | Variable Description(s) | Formula |
| --- | --- | --- |
| A. geoMean of each measurement, calculated from the modern samples | $GM_x$ where $x$ = measurement type (ex: FHH) | $\left( \prod_{i=1}^n GM_{x_i} \right)^{\frac{1}{n}} = \sqrt[n]{x_{y1} \cdot x_{y2} \cdot \dots \cdot x_{yn}}$ |
| B. For each individual bone and element, fold change from geoMean | $FC_{xy}$ where $y$ = individual (ex: BMOC001) | $FC_{xy1} = \left( \frac{x_{y1}}{GM_x} \right) - 1$ |
| C. Average fold change for each skeletal element | $A_{z1y1}$ where $z$ = sided element (ex: right femur) | $A_{z1y1} = \frac{FC_{x1y1} + \dots + FC_{xn_{y1}}}{n}$ |
| D. Aggregate per-individual fold-change scores from all available skeletal elements for that individual | $AS_{y1}$ | $AS_{y1} = \frac{A_{z1y1} + A_{z2y1} + \dots + A_{zn_{y1}}}{n}$ |

Then, all of the FC values available for each separate sided bone (*z*) were arithmetically averaged (*A*; **Table 3, Step C**). Finally, for each modern individual an aggregate score (*AS*) was calculated as the mean of the averages from each element available for that individual (up to four; i.e. right and left, humeri and femora) (**Table 3, Step D**). For the subfossil remains, the per-bone average FC value is the same as the individual *AS*, because potential individual-associations among any of the different subfossil elements are unknown.

As an example, modern individual BMOC-001 had two complete femora and humeri, and all measurements were collected. The fold change difference was calculated for each of BMOC-001’s right femoral measurements, and the average of those fold changes was calculated to be -0.005. A negative average fold change (*A*) indicates that the right femoral bones of BMOC-001 were slightly smaller than the right femoral geoMean of the modern population of Beza Mahafaly lemurs. Each of the *A* values for BMOC-001’s two femora and humeri were averaged for a final aggregate score (*AS*). A positive *AS* of 0.003 indicates that the bones of BMOC-001 were slightly larger than the average AS of the modern population of Beza Mahafaly lemurs. Subfossil element UM-TAO-66-21 had only three measurements taken, and the *AS* of the fold changes for those three measurements was 0.142, indicating an element larger than BMOC-001 as well as the average *AS* of the modern population.

We conducted two randomized subsampling permutation analyses to determine where the subfossil dataset’s average aggregate score would fall against distributions generated from the same amount of modern measurement data. One permutation analysis was conducted with the full n=12 subfossil elements treated as separate individuals and the other by analyzing only the skeletal element and side with the best representation in the dataset, or the minimum number of individuals (right distal femur; MNI n=4). A random subset of the modern data was partitioned to mimic those available for the subfossil individuals. In the case of the MNI group, this meant two right femoral FBE measurements and four DML2 measurements. The *GM, FC*, and *AS* were calculated from this subset of modern measurements, as described above, and then the entire subset procedure was repeated 10,000 times for comparison with the average aggregate scores of the respective subfossil groupings. All code developed and used for this project is available in the GitHub repository https://github.com/AlexisPSullivan/Sifaka.

## RESULTS

We compared long bone measurements as a proxy for body size between *Propithecus verreauxi* skeletal remains from the Taolambiby subfossil site (725-560 – 1075-955 calibrated 95.4% CI years before present) to those collected from modern individuals of the same species at the nearby Beza Mahafaly Special Reserve (<10km from Taolambiby) to evaluate whether this population experienced body size diminution in the recent past. If so, then this result would be consistent with the hypothesis that human size-selective hunting pressures may have driven phenotypic evolutionary change in Madagascar’s surviving fauna.

When comparing subfossil and modern specimens, we treated the entire scanned collection (N = 12) as our maximum number of individuals (MAX, **Figure 3A**), and the four subfossil right distal femoral fragments (UM-TAO-66-26, -29, -32, -33) as our minimum number of individuals (MNI, **Figure 4A**). We calculated the fold-change difference between the measurements of each bone (subfossil) or individual (modern) and the geoMean of the modern population (**Figures 3B**; **Figure 4B**).

**Figure 3:**
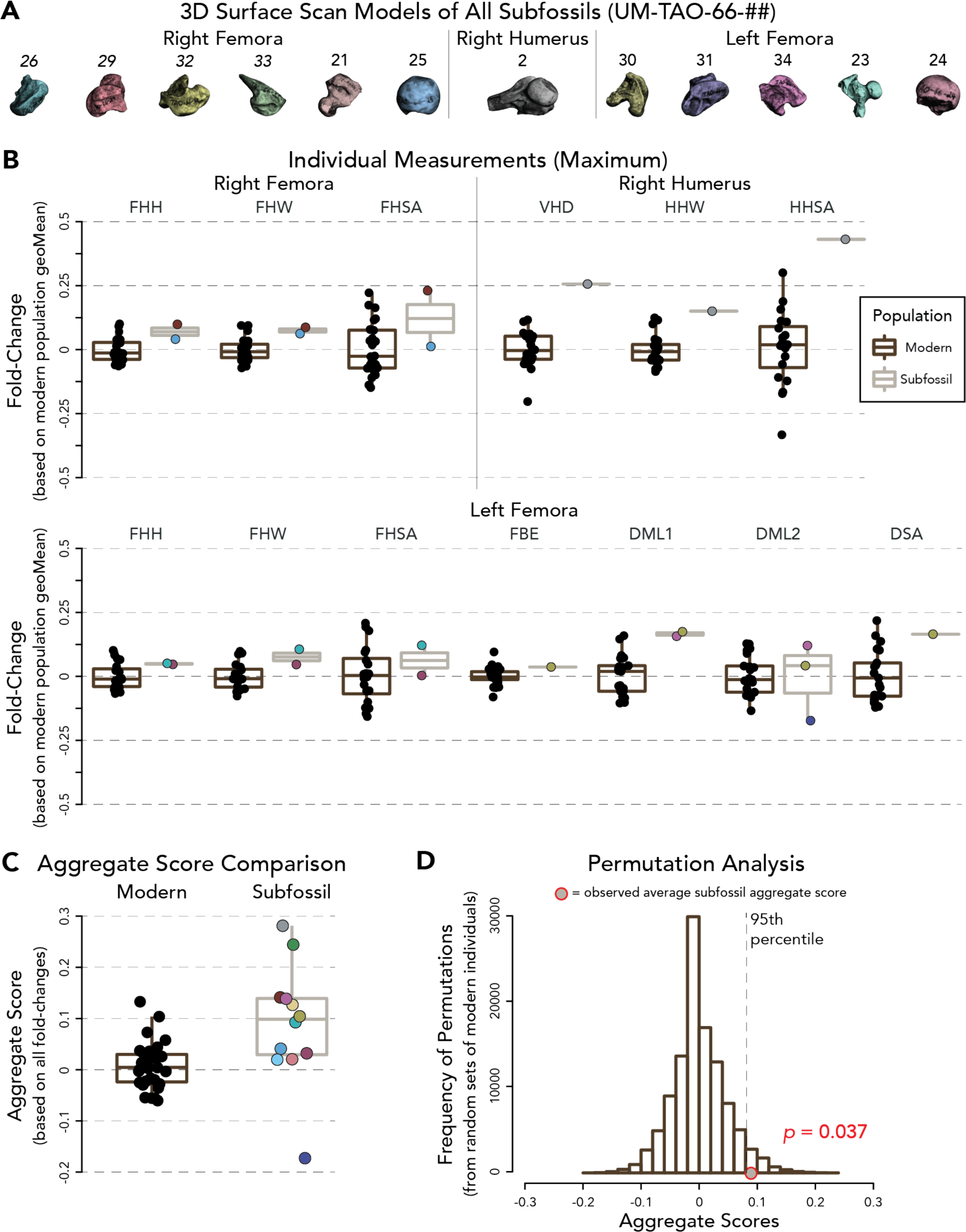
Our comparative morphological results for the maximum number of subfossil individuals (MAX). A: Each subfossil sample is separately colored to demonstrate progression through our pipeline. B: Fold-changes for both modern and subfossil 3D surface scan measurements for which there any subfossil data points are available. See Table 1 for the description of individual measurements. C. Aggregate scores for each subfossil skeletal element and modern individual. D. Permutation analysis depicting the distribution of average aggregate scores calculated from 10,000 subsets of modern measurements randomly selected to match the sample sizes of the MAX subfossil dataset. The actual average aggregate value (0.089) for the MAX subfossil sample is shown with a red circle. The indicated empirical p-value (p=0.037) represents the proportion of permuted modern values equal or greater to the actual subfossil value.

**Figure 4:**
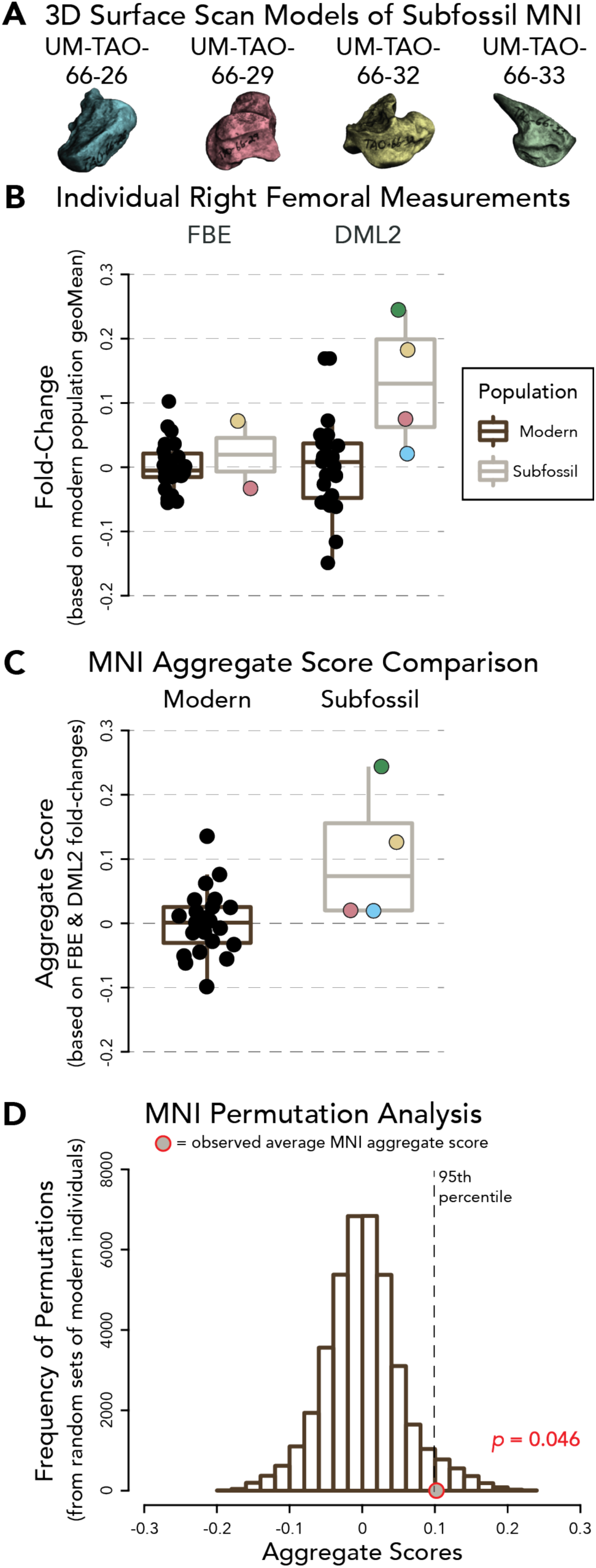
Comparative morphological results for the minimum number of subfossil individuals (MNI). A: Each subfossil sample is separately colored to demonstrate progression through our pipeline. B: Fold-changes for those modern and subfossil 3D surface scan measurements for which there are any subfossil data points. See Table 1 for the description of individual measurements. C. Aggregate scores for each subfossil and modern individual. D. Permutation analysis depicting the distribution of average aggregate scores calculated from 10,000 subsets of modern measurements randomly selected to match the sample sizes of the MNI subfossil dataset. The actual average aggregate value (0.103) for the MNI subfossil sample is shown with a red circle. The indicated empirical p-value (p=0.046) represents the proportion of permuted modern values equal or greater to the actual subfossil value.

As described in the Methods, each of the fold-change averages was arithmetically averaged for each individual across every element available for that individual to create an aggregate score (AS) that we used to directly compare modern and subfossil individuals to each other (**Figure 3C; Figure 4C; Supplementary Table 6**). The mean aggregate score for all subfossil elements (MAX; 0.089±0.117) is significantly greater than that for the modern individuals (0.009±0.045; Welch two-sample t-test; p=0.039; **Figure 3C**). With only four right distal subfossil femora, the mean aggregate score for subfossil MNI (0.103±0.107) was not significantly different than that for the modern individuals with a t-test (0.002±0.051; p=0.153; **Figure 4C**).

We further used a permutation scheme to test the null hypothesis of no size difference between the modern and subfossil populations. Specifically, for each of the MAX and MNI comparisons we selected a random subset of the modern data to match the number of specimens and measurement types of the subfossil dataset, computed the aggregate score for that permuted modern dataset, and repeated that process 10,000 times. To compute empirical p-values, the observed subfossil aggregate scores were compared to the distributions of permuted results, with significant differences for both MAX (**Figure 3D**; p=0.037) and MNI (**Figure 4D**; p=0.046).

## DISCUSSION

This work represents the first systematic assessment of the potential evolutionary effects of human size-selective hunting pressures on body size in a non-human primate. Using skeletal remains of both modern and subfossil *P. verreauxi* individuals from the same region of Southwest Madagascar and a high-resolution 3D surface scanning-based approach, we found that archaeological (725-560 – 1075-955 cal. years before present) body size-associated skeletal measurements were significantly larger than those of the modern sample.

While this result is consistent with the hypothesis of recent phyletic dwarfism in response to size-selective hunting pressures by humans, our finding alone does not necessarily demonstrate a history of adaptive evolution for smaller body sizes in this population. As an alternative explanation, the archaeological sample could be biased by assemblage and/or taphonomic processes [48]. For example, if past people were preferentially hunting larger sifaka with projectiles such as slingshots or blowguns [49,50], then individuals who ended up in the archaeological sample may have been larger than the average for the overall population at the time. Additionally, larger bones may have been more likely to be preserved in the Taolambiby wash. The combination of expanded archaeological sampling and evolutionary genomic analyses with knowledge of *P. verreauxi* body size-associated alleles may ultimately be necessary to distinguish between the evolutionary vs. assemblage/taphonomic scenarios.

If the inferred body size difference between the archaeological and modern *P. verreauxi* samples does reflect an evolutionary process in response to human size-selective hunting behavior, then it is of interest to compare the estimated rate of that evolutionary change to previously observations for other archaeological cases of this phenomenon. We estimated an evolutionary rate of 156 *darwins*, or the magnitude of morphological change (absolute value of the difference between the natural log of the starting trait value and the natural log of the ending trait value) per million years [51] using our data from the right femur width of lateral distal condyle (n=4 subfossil individuals, mean width=7.76 mm, average 771 cal. years BP, calculated with midpoints of 95.4% CI cal. years BP; n=23 modern individuals, mean width=6.88 mm). Using these values along with the *P. verreauxi* cohort generation time (average between the birth of a female and the birth of her daughters) of 18.5 years [52,53], we also estimated a *Haldane (H)* evolutionary rate of -0.039 *H*, which represents the rate of change in standard deviations per generation [54]. While *H* is likely a more meaningful statistic, only *darwin* estimates are available for direct comparison with other taxa. Compared to seven previously-analyzed archaeological examples of purported morphological change in response to human behavior (*darwin* mean=21.4; range = 1-72; [1] the rate of 156 *darwins* for *P. verreauxi* is at least ∼2x and an average of 7.3x greater.

## ACKNOWLEDGMENTS

We thank the Madagascar National Parks organization and our colleagues at the Beza Mahafaly Special Reserve, including Joel Ratsirarson, Jeannin Ranaivonasy, Sibien Mahereza, Elaine Guevara, Brenda Bradley, Roshna Wunderlich, and Alison Richard. We also thank the University of Massachusetts Amherst Natural History Collections for allowing access to the Taolambiby collection, and Lily Doershuk for her advice on working with 3D data. We would like to acknowledge support from the Peter A. and Marion W. Schwartz Family Foundation for providing funds (to L.R.G.) to help build the Beza Mahafaly Osteological Collection. This project was supported by National Science Foundation grant BCS-1554834 (to G.H.P.), the National Science Foundation grant BCS-1750598 (to L.R.G.), and the National Science Foundation Graduate Research Fellowship Program (DGE1255832, to A.P.S.). Any opinions, findings, and conclusions or recommendations expressed in this material are those of the authors and do not necessarily reflect the views of the National Science Foundation.

## SUPPLEMENTARY MATERIALS

**Supplemental Database:** MorphoSource repository of Artec Space Spider 3D surface scan data (.ply models) for all modern and subfossil *Propithecus verreauxi* individuals (https://www.morphosource.org/Detail/ProjectDetail/Show/project_id/698)

**Supplemental Table 1**: List of modern Beza Mahafaly Propithecus verreauxi individuals and measured long bones (femur, humerus)

**Supplemental Table 2**: Modern and subfossil *Propithecus verreauxi* radial caliper measurements and fold-changes

**Supplemental Table 3**: Modern and subfossil *Propithecus verreauxi* 3D scan Avizo measurements and fold-changes

**Supplemental Table 4:** Average percent differences between radial caliper and 3D model measurements

**Supplemental Table 5**: Radial caliper versus 3D scan Avizo measurements

**Supplemental Table 6**: Aggregate scores for all modern and subfossil *Propithecus verreauxi* individuals

