## Supplementary Materials for "Potential evolutionary body size reduction in a Malagasy primate (*Propithecus verreauxi*) in response to human size-selective hunting pressure"

**Supplemental Database:** MorphoSource repository of Artec Space Spider 3D surface scan data (.ply models) for all modern and subfossil *Propithecus verreauxi* individuals (https://www.morphosource.org/Detail/ProjectDetail/Show/project_id/698)

**Supplemental Table 1**: List of modern Beza Mahafaly Propithecus verreauxi individuals and measured long bones (femur, humerus)

**Supplemental Table 2**: Modern and subfossil *Propithecus verreauxi* radial caliper measurements and fold-changes

**Supplemental Table 3**: Modern and subfossil *Propithecus verreauxi* 3D scan Avizo measurements and fold-changes

**Supplemental Table 4:** Average percent differences between radial caliper and 3D model measurements

**Supplemental Table 5**: Radial caliper versus 3D scan Avizo measurements

**Supplemental Table 6**: Aggregate scores for all modern and subfossil *Propithecus verreauxi* individuals

**Supplementary Table 1: List of Modern Beza Mahafaly Propithecus verreauxi Individuals and Measured Long Bones (Femur, Humerus)**

| Drawer #* | Sample ID # | # Scanned Elements | # Scanned Femora | # Scanned Humeri | Date Recovered | Finder |
|---|---|---|---|---|---|---|
| 1 | BMOC004 | 5 | 2 | 3 (right broken into two pieces) | 7/17/93 | Kashka Kubzdela |
| 1 | BMOC125 | 1 | 0 | 1 (right) | 6/10/03 | Krista Fish |
| 1 | BMOC008 | 4 | 2 | 2 | 1/27/94 | Kashka Kubzdela |
| 1 | BMOC019 | 4 | 2 | 2 | | Esoes |
| 1 | BMOC172 | 4 | 2 | 2 | 3/31/08 | Enafa |
| 1 | BMOC030 | 2 | 1 (right) | 1 (right) | 8/1/01 | |
| 1 | BMOC001 | 4 | 2 | 2 | 7/23/93 | Kashka Kubzdela |
| 1 | BMOC015 | 4 | 2 | 2 | 7/11/94 | Kashka Kubzdela |
| 1 | BMOC073 | 4 | 2 | 2 | 8/4/01 | Sifaka monitoring team (Diane Brockman) |
| 1 | BMOC043 | 2 | 0 | 2 | | |
| 1 | BMOC020 | 4 | 2 | 2 | 10/25/94 | Esoahere and Esoes |
| 2 | BMOC035 | 1 | 1 (right) | 0 | | |
| 2 | BMOC169 | 1 | 1 (right) | 0 | | |
| 2 | Fogel 2012 | 2 | 0 | 2 | 2/11/12 | Andy Fogel |
| 2 | BMOC174 | 4 | 2 | 2 | 1/17/09 | Enafa |
| 2 | BMOC005 | 2 | 2 | 0 | 8/12/94 | Enafa and Kashka Kubzdela |
| 2 | BMOC075 | 4 | 2 | 2 | 8/-/2001 | Julie Parks |
| 2 | BMOC021 | 2 | 2 | 0 | 1/11/95 | Zavie |
| 2 | BMOC156 | 1 | 1 (right) | 0 | 6/30/06 | |
| 2 | BMOC197 | 4 | 2 | 2 | 6/28/06 | Anne Axel and Max |
| 2 | BMOC157 | 4 | 2 | 2 | 7/3/06 | |
| 2 | BMOC014 | 2 | 2 | 0 | 6/21/94 | Anje Van Berckelaer |
| 2 | BMOC142 | 4 | 2 | 2 | 8/1/05 | Diane Brockman |
| 2 | BMOC180 | 3 | 1 (left) | 2 | 11/27/07 | Efitria |
| 2 | BMOC028 | 4 | 2 | 2 | | |
| 2 | BMOC191 | 1 | 1 (left) | 0 | After 9/2008 | Efitria |
| 2 | BMOC163 | 1 | 1 (left) | 0 | 6/29/06 | Anne Axel |
| 2 | BMOC173 | 4 | 2 | 2 | | Enafa |
| 2 | BMOC137 | 4 | 2 | 2 | 7/17/05 | Anne Axel, E. Rasoazanabary, L. Godfrey |
| 5 | Indiv. #570 | 4 | 2 | 2 | 6/-/2012 | Katie Grogan and Tahiri |
| 5 | Indiv. #550 | 4 | 2 | 2 | 5/-/2008 | |
| *Top->bottom | | 94 | 49 (25 right, 24 left) | 45 (23 right, 21 left) | | |

**Contiued**

| Drawer #* | Sample ID # | Collar Number/FN ID # | Sex | Location | Other Recorded Information |
|---|---|---|---|---|---|
| 1 | BMOC004 | | | 20-30m West of Blue W/4m North of the trail South of Blue 1 | Black collar |
| 1 | BMOC125 | | | Blue 1 between Orange W and Pink W but closer to Orange W | |
| 1 | BMOC008 | | | 10m East of Blue W/3m North of Blue 2 | |
| 1 | BMOC019 | 40 | | 3m West of Red W/45m North of Pink 3 | Group Zavma1 |
| 1 | BMOC172 | 549 | Male | | Last seen 2/2008, group Saksud |
| 1 | BMOC030 | | | North of Pink 1/East of Orange W | |
| 1 | BMOC001 | 242 | | 20m East of Yellow W/Pink 2 | |
| 1 | BMOC015 | 204 | Female | 2m East of Yellow W/30m North of Blue 3 | Found 5/12/1994 (dead ~16 days), group Lavaka, born 1981 |
| 1 | BMOC073 | 156 | | South of Blue 2/East of center | |
| 1 | BMOC043 | | | | |
| 1 | BMOC020 | 26 | | 50m South of Pink 2/8m East of Yellow W | Sifaka mate nimiro |
| 2 | BMOC035 | | | | |
| 2 | BMOC169 | | | | |
| 2 | Fogel 2012 | | | Orange E - river trail, Pink 2 - Pink3 (23 39.041 S, 44 58.126 E) | |
| 2 | BMOC174 | 288 | Male | South Pink 2/West of Orange W | Indiv. # 288 (96), Borety; last seen 1/2007, discovered on a tree (7/2007), mort par un diarhee,<br>buried by Bob Dewar (2007); broken left humerus |
| 2 | BMOC005 | 74 | Female | 50m West of Black/20m South of Pink 1 | Group Nify, born 1987, died 8/1993 |
| 2 | BMOC075 | 224 | | Just West of center/North of Pink 2 | |
| 2 | BMOC021 | | | 30m North of Pink/25m West of Green W | |
| 2 | BMOC156 | | | | Designated "Prop E"; WGS 84, UTM 38S, 460704, 7383447; rat damage |
| 2 | BMOC197 | | | Within Parcel 1, near northern boundary of res. | Designated "Prop H 2006", grey collar, no tag; UTM 38S, WGS 84, 0462526, 7384463;<br>older individual, skull (cranium and mandible), assorted postcrania including ilium, ischium, femora, ribs, tibia, 1 ulna, 1 clavicle, 1 scapula |
| 2 | BMOC157 | | | | Designated "Prop F"; UTM 38S, WGS 84, 462571, 7384986;<br>assorted post crania, *Kristi Lewton - 3 tibia in bag |
| 2 | BMOC014 | | | 25m West of Pink E/50m North of Blue 2 | Found with dry stomach contents |
| 2 | BMOC142 | 9093 | Male | ~10m South of Pink 2, 3m East of Orange W | Complete skeleton, 11 years old, group Borety |
| 2 | BMOC180 | | | Pink 4 5m from Red | Skull and mandible (complete), post crania bones |
| 2 | BMOC028 | 179 | | | |
| 2 | BMOC191 | 764 | Male | Parcel 1 | Last seen 9/2008, NB: no data, skull, mandible, femur |
| 2 | BMOC163 | | | | Designated "Prop L 2006"; UTM 38S, WGS 84, 0461187, 7383361;<br>very worn teeth, scavenger damage, cranium and mandible, assorted postcranial material including 1 tibia, femora, left and right innominate, 2 ulna, 1 clavicle, sacrum, 1 partial fibula, misc. vertebrae and phallanges |
| 2 | BMOC173 | 367 | Female | North of Pink 1/East of Yellow E | Born 1998, died 7/2008; group Vavigoa, Jonarisoa and Jadry |
| 2 | BMOC137 | 282/8 | | Between Pink 2 and 3, close to Green W and Blue W | |
| 5 | Indiv. #570 | | | 100m West of Beza camp, across N/S side road | |
| 5 | Indiv. #550 | | Male | Nord vala nord (13m) SW la piste de Mahazarivo (red) | Group NW1GPMAIO8, Nico group, Mont de la janvier 2009, Ralaivao |

**Supplementary Table 2: Modern and subfossil Propithecus verreauxi radial caliper measurements and fold-changes**

| | | | | | *RAW MEASUREMENT (mm)* | | | | | | | | *FOLD-CHANGE (from modern geoMean (mm))* | | | | |
|---|---|---|---|---|---|---|---|---|---|---|---|---|---|---|---|---|---|
| *SIDE* | *ELEMENT* | *AGE* | *COLLECTION* | *SAMPLE ID* | *MFL* | *MFD* | *MFC* | *FHH* | *FHW* | *FBE* | *L (178.3* | *FD (8.77* | *MFC (29.06)* | *FHH (12.00)* | *FHW (12.21)* | *FBE (20.12)* | *CHANGE (SIDED)* |
| Right | Femur | Modern | BMOC | 550 | 169.34 | 8.59 | 28.14 | 11.71 | 12 | 18.96 | -0.05 | -0.02 | -0.03 | -0.02 | -0.02 | -0.06 | -0.03 |
| Right | Femur | Modern | BMOC | 570 | 172.72 | 9.05 | 29.47 | 11.66 | 11.84 | 19.46 | -0.03 | 0.03 | 0.01 | -0.03 | -0.03 | -0.03 | -0.01 |
| Right | Femur | Modern | BMOC | BMOC001 | 187.4 | 8.24 | 29.02 | 11.92 | 12.2 | 20.28 | 0.05 | -0.06 | 0 | -0.01 | 0 | 0.01 | 0 |
| Right | Femur | Modern | BMOC | BMOC004 | 173.9 | 8.51 | 29.79 | 11.5 | 11.82 | 20.06 | -0.03 | -0.03 | 0.03 | -0.04 | -0.03 | 0 | -0.02 |
| Right | Femur | Modern | BMOC | BMOC005 | 185.44 | 9.13 | 30.48 | 12.53 | 12.93 | 20.54 | 0.04 | 0.04 | 0.05 | 0.04 | 0.06 | 0.02 | 0.04 |
| Right | Femur | Modern | BMOC | BMOC008 | 187.04 | 8.73 | 29.88 | 13.19 | 13.74 | 20.71 | 0.05 | 0 | 0.03 | 0.1 | 0.13 | 0.03 | 0.05 |
| Right | Femur | Modern | BMOC | BMOC014 | 172.19 | 8.62 | 29.19 | 11.94 | 12.43 | 20.54 | -0.03 | -0.02 | 0 | 0 | 0.02 | 0.02 | 0 |
| Right | Femur | Modern | BMOC | BMOC015 | 166.48 | 8.22 | 27.88 | 12.7 | 12.56 | 19.08 | -0.07 | -0.06 | -0.04 | 0.06 | 0.03 | -0.05 | -0.02 |
| Right | Femur | Modern | BMOC | BMOC019 | 180.74 | 8.89 | 29.98 | 12.6 | 12.27 | 21.23 | 0.01 | 0.01 | 0.03 | 0.05 | 0 | 0.06 | 0.03 |
| Right | Femur | Modern | BMOC | BMOC020 | 161.88 | 8.72 | 29.9 | 11.81 | 12.07 | 19.61 | -0.09 | -0.01 | 0.03 | -0.02 | -0.01 | -0.03 | -0.02 |
| Right | Femur | Modern | BMOC | BMOC021 | 195.32 | 8.75 | 29.54 | 12.13 | 12.27 | 19.71 | 0.09 | 0 | 0.02 | 0.01 | 0 | -0.02 | 0.02 |
| Right | Femur | Modern | BMOC | BMOC028 | NA | NA | NA | NA | NA | 20.06 | NA | NA | NA | NA | NA | 0 | 0 |
| Right | Femur | Modern | BMOC | BMOC030 | NA | 9.26 | 30.47 | 12.34 | 12.82 | NA | NA | 0.06 | 0.05 | 0.03 | 0.05 | NA | 0.05 |
| Right | Femur | Modern | BMOC | BMOC035 | 178.4 | 8.96 | 30.52 | 11.81 | 11.99 | 19.45 | 0 | 0.02 | 0.05 | -0.02 | -0.02 | -0.03 | 0 |
| Right | Femur | Modern | BMOC | BMOC073 | 174.16 | 8.52 | 28.54 | 12.64 | 12.71 | 19.91 | -0.02 | -0.03 | -0.02 | 0.05 | 0.04 | -0.01 | 0 |
| Right | Femur | Modern | BMOC | BMOC075 | NA | 8.33 | 27.58 | 11.95 | 12.45 | 20.03 | NA | -0.05 | -0.05 | 0 | 0.02 | 0 | -0.02 |
| Right | Femur | Modern | BMOC | BMOC137 | 186.57 | 9.04 | 31.26 | 13.21 | 13.65 | 21.93 | 0.05 | 0.03 | 0.08 | 0.1 | 0.12 | 0.09 | 0.08 |
| Right | Femur | Modern | BMOC | BMOC142 | 176.94 | 9.05 | 29.71 | 13.19 | 13.3 | 20.06 | -0.01 | 0.03 | 0.02 | 0.1 | 0.09 | 0 | 0.04 |
| Right | Femur | Modern | BMOC | BMOC156 | 180.63 | 9.9 | 30.49 | 12.12 | 11.73 | 20.33 | 0.01 | 0.13 | 0.05 | 0.01 | -0.04 | 0.01 | 0.03 |
| Right | Femur | Modern | BMOC | BMOC157 | 188.36 | 8.89 | 29.46 | 12.39 | 12.91 | 19.84 | 0.06 | 0.01 | 0.01 | 0.03 | 0.06 | -0.01 | 0.03 |
| Right | Femur | Modern | BMOC | BMOC169 | 179.27 | 8.89 | 28.42 | 8.05 | 8.26 | 21.5 | 0.01 | 0.01 | -0.02 | -0.33 | -0.32 | 0.07 | -0.1 |
| Right | Femur | Modern | BMOC | BMOC172 | 168.99 | 8.32 | 25.36 | 11.88 | 12 | 19.26 | -0.05 | -0.05 | -0.13 | -0.01 | -0.02 | -0.04 | -0.05 |
| Right | Femur | Modern | BMOC | BMOC173 | 181.36 | 8.34 | 27.59 | 12.03 | 12.28 | 19.84 | 0.02 | -0.05 | -0.05 | 0 | 0.01 | -0.01 | -0.01 |
| Right | Femur | Modern | BMOC | BMOC174 | 182.7 | 8.86 | 28.05 | 12.12 | 12.26 | 20.66 | 0.02 | 0.01 | -0.03 | 0.01 | 0 | 0.03 | 0.01 |
| Right | Femur | Modern | BMOC | BMOC197 | NA | 8.77 | 27.36 | 11.65 | 11.68 | NA | NA | 0 | -0.06 | -0.03 | -0.04 | NA | -0.03 |
| Right | Femur | Subfossil | TAO | TAO-66-21 | NA | NA | NA | 13.86 | 13.62 | NA | NA | NA | NA | 0.16 | 0.12 | NA | 0.14 |
| Right | Femur | Subfossil | TAO | TAO-66-25 | NA | NA | NA | 13.21 | 13.18 | NA | NA | NA | NA | 0.1 | 0.08 | NA | 0.09 |
| Right | Femur | Subfossil | TAO | TAO-66-32 | NA | NA | NA | NA | NA | 21.48 | NA | NA | NA | NA | NA | 0.07 | 0.07 |

| | | | | | *RAW MEASUREMENT (mm)* | | | | | | | | *FOLD-CHANGE (from modern geoMean (mm))* | | | | |
|---|---|---|---|---|---|---|---|---|---|---|---|---|---|---|---|---|---|
| *SIDE* | *ELEMENT* | *AGE* | *COLLECTION* | *SAMPLE ID* | *MFL* | *MFD* | *MFC* | *FHH* | *FHW* | *FBE* | *L (177.1* | *FD (8.66* | *MFC (29.00)* | *FHH (12.33)* | *FHW (12.54)* | *FBE (20.09)* | *CHANGE (SIDED)* |
| Left | Femur | Modern | BMOC | 550 | 167.34 | 8.24 | 28.31 | 11.83 | 12.04 | 19.41 | -0.05 | -0.05 | -0.02 | -0.04 | -0.04 | -0.03 | -0.04 |
| Left | Femur | Modern | BMOC | 570 | 174.18 | 9.04 | 29.94 | 11.74 | 11.94 | 19.47 | -0.02 | 0.04 | 0.03 | -0.05 | -0.05 | -0.03 | -0.01 |
| Left | Femur | Modern | BMOC | BMOC001 | 187.64 | 8.05 | 27.99 | 12.14 | 12.28 | 20.02 | 0.06 | -0.07 | -0.03 | -0.02 | -0.02 | 0 | -0.01 |
| Left | Femur | Modern | BMOC | BMOC004 | 174.3 | 8.9 | 29.2 | 11.51 | 11.83 | 20.12 | -0.02 | 0.03 | 0.01 | -0.07 | -0.06 | 0 | -0.02 |
| Left | Femur | Modern | BMOC | BMOC005 | NA | 9.1 | 30.09 | NA | NA | 20.42 | NA | 0.05 | 0.04 | NA | NA | 0.02 | 0.03 |
| Left | Femur | Modern | BMOC | BMOC008 | 187.63 | 8.5 | 29.83 | 13.39 | 13.57 | 20.64 | 0.06 | -0.02 | 0.03 | 0.09 | 0.08 | 0.03 | 0.04 |
| Left | Femur | Modern | BMOC | BMOC014 | NA | 8.22 | 23.8 | NA | NA | 20.72 | NA | -0.05 | -0.18 | NA | NA | 0.03 | -0.07 |
| Left | Femur | Modern | BMOC | BMOC015 | 165.93 | 8.4 | 27.49 | 12.8 | 12.72 | 18.98 | -0.06 | -0.03 | -0.05 | 0.04 | 0.01 | -0.06 | -0.02 |
| Left | Femur | Modern | BMOC | BMOC019 | NA | 8.85 | 31.55 | NA | NA | NA | NA | 0.02 | 0.09 | NA | NA | NA | 0.05 |
| Left | Femur | Modern | BMOC | BMOC020 | 167.05 | 8.7 | 29.62 | 11.97 | 12.22 | 19.84 | -0.06 | 0 | 0.02 | -0.03 | -0.03 | -0.01 | -0.02 |
| Left | Femur | Modern | BMOC | BMOC021 | 174.93 | 8.62 | 30.02 | 12.23 | 12.37 | 19.56 | -0.01 | 0 | 0.04 | -0.01 | -0.01 | -0.03 | 0 |
| Left | Femur | Modern | BMOC | BMOC028 | 185.8 | 8.54 | 30.5 | 12.52 | 12.7 | 20.36 | 0.05 | -0.01 | 0.05 | 0.02 | 0.01 | 0.01 | 0.02 |
| Left | Femur | Modern | BMOC | BMOC073 | 174.65 | 8.62 | 28.45 | 12.2 | 12.55 | 19.76 | -0.01 | 0 | -0.02 | -0.01 | 0 | -0.02 | -0.01 |
| Left | Femur | Modern | BMOC | BMOC075 | 171.54 | 8.29 | 27.88 | 12.09 | 12.47 | 20.12 | -0.03 | -0.04 | -0.04 | -0.02 | -0.01 | 0 | -0.02 |
| Left | Femur | Modern | BMOC | BMOC137 | 187.06 | 9.28 | 31.66 | 13.41 | 13.56 | 21.76 | 0.06 | 0.07 | 0.09 | 0.09 | 0.08 | 0.08 | 0.08 |
| Left | Femur | Modern | BMOC | BMOC142 | 177.34 | 8.78 | 29.02 | 13.1 | 13.21 | 20.25 | 0 | 0.01 | 0 | 0.06 | 0.05 | 0.01 | 0.02 |
| Left | Femur | Modern | BMOC | BMOC157 | 189.02 | 8.63 | 29.9 | 12.4 | 12.84 | 20.73 | 0.07 | 0 | 0.03 | 0.01 | 0.02 | 0.03 | 0.03 |
| Left | Femur | Modern | BMOC | BMOC163 | NA | 9.42 | 32.87 | 13.37 | 13.66 | NA | NA | 0.09 | 0.13 | 0.08 | 0.09 | NA | 0.1 |
| Left | Femur | Modern | BMOC | BMOC172 | 168.54 | 8.55 | 26.75 | 11.95 | 11.88 | 18.88 | -0.05 | -0.01 | -0.08 | -0.03 | -0.05 | -0.06 | -0.05 |
| Left | Femur | Modern | BMOC | BMOC173 | 180.75 | 8.31 | 27.23 | 12.18 | 12.34 | 20.21 | 0.02 | -0.04 | -0.06 | -0.01 | -0.02 | 0.01 | -0.02 |
| Left | Femur | Modern | BMOC | BMOC174 | 183.16 | 8.9 | 28.72 | 12.35 | 12.39 | 20.61 | 0.03 | 0.03 | -0.01 | 0 | -0.01 | 0.03 | 0.01 |
| Left | Femur | Modern | BMOC | BMOC180 | NA | 9.04 | 29.95 | 12.02 | 12.82 | NA | NA | 0.04 | 0.03 | -0.03 | 0.02 | NA | 0.02 |
| Left | Femur | Modern | BMOC | BMOC191 | NA | 8.55 | 28.21 | NA | NA | 20.56 | NA | -0.01 | -0.03 | NA | NA | 0.02 | -0.01 |
| Left | Femur | Modern | BMOC | BMOC197 | 172.43 | 8.51 | 28.48 | 11.7 | 11.69 | 19.73 | -0.03 | -0.02 | -0.02 | -0.05 | -0.07 | -0.02 | -0.03 |
| Left | Femur | Subfossil | TAO | TAO-66-23 | NA | NA | NA | 13.06 | 13.52 | NA | NA | NA | NA | 0.06 | 0.08 | NA | 0.07 |
| Left | Femur | Subfossil | TAO | TAO-66-24 | NA | NA | NA | 13.24 | 13.1 | NA | NA | NA | NA | 0.07 | 0.04 | NA | 0.06 |
| Left | Femur | Subfossil | TAO | TAO-66-30 | NA | NA | NA | NA | NA | 21.23 | NA | NA | NA | NA | NA | 0.06 | 0.06 |
| Left | Femur | Subfossil | TAO | TAO-66-34 | NA | NA | NA | NA | NA | 20.99 | NA | NA | NA | NA | NA | 0.04 | 0.04 |

| | | | | | *RAW MEASUREMENT (mm)* | | | | | *FOLD-CHANGE (from modern geoMean (mm))* | | | | | |
|---|---|---|---|---|---|---|---|---|---|---|---|---|---|---|---|
| *SIDE* | *ELEMENT* | *AGE* | *COLLECTION* | *SAMPLE ID* | *MHL* | *MHD* | *VHD* | *HHW* | *BBH* | *HL (91.1* | *HD1 (6.6* | *HD (11.17* | *HHW (9.61)* | *BBH (20.88)* | *CHANGE (SIDED)* |
| Right | Humerus | Modern | BMOC | 550 | 88.55 | 6.42 | 10.89 | 9.26 | 20.41 | -0.03 | -0.04 | -0.02 | -0.04 | -0.02 | -0.03 |
| Right | Humerus | Modern | BMOC | 570 | 89.66 | 6.18 | 10.81 | 9 | 19.76 | -0.02 | -0.07 | -0.03 | -0.06 | -0.05 | -0.05 |
| Right | Humerus | Modern | BMOC | BMOC001 | 94.53 | 6.32 | 10.95 | 9.63 | 21.45 | 0.04 | -0.05 | -0.02 | 0 | 0.03 | 0 |
| Right | Humerus | Modern | BMOC | BMOC004 | NA | 6.83 | 10.14 | 9.02 | 20.97 | NA | 0.03 | -0.09 | -0.06 | 0 | -0.03 |
| Right | Humerus | Modern | BMOC | BMOC008 | 94.63 | 7.14 | 12.2 | 10.44 | NA | 0.04 | 0.07 | 0.09 | 0.09 | NA | 0.07 |
| Right | Humerus | Modern | BMOC | BMOC015 | 86.77 | 7.18 | 11.49 | 10.13 | 21.31 | -0.05 | 0.08 | 0.03 | 0.05 | 0.02 | 0.03 |
| Right | Humerus | Modern | BMOC | BMOC019 | 92.68 | 7.16 | 11.4 | 10.24 | 21.49 | 0.02 | 0.08 | 0.02 | 0.07 | 0.03 | 0.04 |
| Right | Humerus | Modern | BMOC | BMOC020 | 88.86 | 7.07 | 10.67 | 9.79 | 20.91 | -0.03 | 0.06 | -0.04 | 0.02 | 0 | 0 |
| Right | Humerus | Modern | BMOC | BMOC028 | 92.49 | 5.53 | 12.06 | 9.39 | 20.85 | 0.01 | -0.17 | 0.08 | -0.02 | 0 | -0.02 |
| Right | Humerus | Modern | BMOC | BMOC030 | NA | 7.6 | 10.95 | 9.63 | NA | NA | 0.14 | -0.02 | 0 | NA | 0.04 |
| Right | Humerus | Modern | BMOC | BMOC043 | 89.54 | 7.13 | 10.67 | 9.39 | 20.98 | -0.02 | 0.07 | -0.04 | -0.02 | 0 | 0 |
| Right | Humerus | Modern | BMOC | BMOC073 | 89.31 | 6.59 | 11.15 | 9.63 | 21.19 | -0.02 | -0.01 | 0 | 0 | 0.01 | 0 |
| Right | Humerus | Modern | BMOC | BMOC075 | NA | 7.32 | NA | NA | 20.14 | NA | 0.1 | NA | NA | -0.04 | 0.03 |
| Right | Humerus | Modern | BMOC | BMOC125 | 97.84 | 6.4 | 11.51 | 10.91 | 22.61 | 0.07 | -0.04 | 0.03 | 0.14 | 0.08 | 0.06 |
| Right | Humerus | Modern | BMOC | BMOC137 | 96.98 | 6.93 | 12.45 | 10.69 | 22.8 | 0.06 | 0.04 | 0.12 | 0.11 | 0.09 | 0.08 |
| Right | Humerus | Modern | BMOC | BMOC142 | 91.9 | 6.49 | 12.11 | 9.86 | 21.52 | 0.01 | -0.03 | 0.08 | 0.03 | 0.03 | 0.02 |
| Right | Humerus | Modern | BMOC | BMOC157 | NA | 6.41 | NA | NA | 20.49 | NA | -0.04 | NA | NA | -0.02 | -0.03 |
| Right | Humerus | Modern | BMOC | BMOC172 | 85.33 | 6.22 | 9.59 | 9.69 | 19.69 | -0.06 | -0.07 | -0.14 | 0.01 | -0.06 | -0.06 |
| Right | Humerus | Modern | BMOC | BMOC173 | 91.27 | 6.35 | 11.26 | 9.01 | 20.66 | 0 | -0.05 | 0.01 | -0.06 | -0.01 | -0.02 |
| Right | Humerus | Modern | BMOC | BMOC174 | 92.53 | 6.42 | 11.55 | 9.04 | 21.11 | 0.02 | -0.04 | 0.03 | -0.06 | 0.01 | -0.01 |
| Right | Humerus | Modern | BMOC | BMOC180 | NA | 6.73 | NA | NA | 20.84 | NA | 0.01 | NA | NA | 0 | 0 |
| Right | Humerus | Modern | BMOC | BMOC197 | 90.05 | 6.81 | 10.88 | 8.81 | 20.23 | -0.01 | 0.02 | -0.03 | -0.08 | -0.03 | -0.03 |
| Right | Humerus | Modern | BMOC | Fogel 2012 | 88.64 | 6.29 | 11 | 9 | 19.35 | -0.03 | -0.06 | -0.01 | -0.06 | -0.07 | -0.05 |
| Right | Humerus | Subfossil | TAO | TAO-66-2 | NA | NA | 14.1 | 10.78 | NA | NA | NA | 0.26 | 0.12 | NA | 0.19 |

| | | | | | *RAW MEASUREMENT (mm)* | | | | | *FOLD-CHANGE (from modern geoMean (mm))* | | | | | |
|---|---|---|---|---|---|---|---|---|---|---|---|---|---|---|---|
| *SIDE* | *ELEMENT* | *AGE* | *COLLECTION* | *SAMPLE ID* | *MHL* | *MHD* | *VHD* | *HHW* | *BBH* | *HL (90.7* | *HD1 (7.1* | *HD (11.37* | *HHW (9.55)* | *BBH (20.77)* | *CHANGE (SIDED)* |
| Left | Humerus | Modern | BMOC | 550 | 88.22 | 6.49 | 11.17 | 8.94 | 20.6 | -0.03 | -0.09 | -0.02 | -0.06 | -0.01 | -0.04 |
| Left | Humerus | Modern | BMOC | 570 | 90.1 | 6.63 | 10.7 | 8.74 | 20.15 | -0.01 | -0.07 | -0.06 | -0.08 | -0.03 | -0.05 |
| Left | Humerus | Modern | BMOC | BMOC001 | 94.36 | 6.16 | 11.05 | 9.84 | 21.64 | 0.04 | -0.14 | -0.03 | 0.03 | 0.04 | -0.01 |
| Left | Humerus | Modern | BMOC | BMOC004 | 90.79 | 6.86 | 10.61 | 9.08 | 21.51 | 0 | -0.04 | -0.07 | -0.05 | 0.04 | -0.02 |
| Left | Humerus | Modern | BMOC | BMOC008 | 96 | 6.68 | 12.39 | 10.45 | 21 | 0.06 | -0.06 | 0.09 | 0.09 | 0.01 | 0.04 |
| Left | Humerus | Modern | BMOC | BMOC015 | 86.56 | 7.61 | 11.57 | 9.97 | 21.11 | -0.05 | 0.07 | 0.02 | 0.04 | 0.02 | 0.02 |
| Left | Humerus | Modern | BMOC | BMOC019 | 92.35 | 7.37 | 11.42 | 10.25 | 21.23 | 0.02 | 0.03 | 0 | 0.07 | 0.02 | 0.03 |
| Left | Humerus | Modern | BMOC | BMOC020 | 88.71 | 6.85 | 10.87 | 9.89 | 20.56 | -0.02 | -0.04 | -0.04 | 0.04 | -0.01 | -0.02 |
| Left | Humerus | Modern | BMOC | BMOC028 | 91.98 | 5.41 | 12.08 | 9.57 | 20.75 | 0.01 | -0.24 | 0.06 | 0 | 0 | -0.03 |
| Left | Humerus | Modern | BMOC | BMOC043 | NA | NA | NA | NA | 20.88 | NA | NA | NA | NA | 0.01 | 0.01 |
| Left | Humerus | Modern | BMOC | BMOC073 | 89.22 | 7.37 | 11.79 | 9.53 | 21.2 | -0.02 | 0.03 | 0.04 | 0 | 0.02 | 0.01 |
| Left | Humerus | Modern | BMOC | BMOC075 | 87.43 | 6.32 | 11.6 | 9.41 | 19.96 | -0.04 | -0.06 | 0.02 | -0.01 | -0.04 | -0.03 |
| Left | Humerus | Modern | BMOC | BMOC137 | 96.88 | 6.85 | 12.34 | 10.51 | 22.72 | 0.07 | -0.04 | 0.09 | 0.1 | 0.09 | 0.06 |
| Left | Humerus | Modern | BMOC | BMOC142 | 92.14 | 6.19 | 11.84 | 9.69 | 21.82 | 0.02 | -0.13 | 0.04 | 0.02 | 0.05 | 0 |
| Left | Humerus | Modern | BMOC | BMOC157 | NA | 6.42 | 11.44 | 9.25 | NA | NA | -0.1 | 0.01 | -0.03 | NA | -0.04 |
| Left | Humerus | Modern | BMOC | BMOC172 | 85.86 | 7.64 | 10.8 | 9.76 | 19.06 | -0.05 | 0.07 | -0.05 | 0.02 | -0.08 | -0.02 |
| Left | Humerus | Modern | BMOC | BMOC173 | 90.88 | 6.2 | 11.1 | 9.03 | 20.36 | 0 | -0.13 | -0.02 | -0.05 | -0.02 | -0.05 |
| Left | Humerus | Modern | BMOC | BMOC174 | 89.68 | 6.99 | 11.43 | 9.07 | 20.82 | -0.01 | -0.02 | 0.01 | -0.05 | 0 | -0.01 |
| Left | Humerus | Modern | BMOC | BMOC180 | 92.96 | 7.4 | 11.59 | 10.3 | 20.56 | 0.02 | 0.04 | 0.02 | 0.08 | -0.01 | 0.03 |
| Left | Humerus | Modern | BMOC | BMOC197 | 90.86 | 6.72 | 11.13 | 8.88 | 20.21 | 0 | -0.06 | -0.02 | -0.07 | -0.03 | -0.03 |
| Left | Humerus | Modern | BMOC | Fogel 2012 | 89.44 | 6.3 | 10.7 | 9.05 | 19.52 | -0.01 | -0.12 | -0.06 | -0.05 | -0.06 | -0.06 |

**Supplementary Table 3: Modern and subfossil Propithecus verreauxi 3D scan Avizo measurements and fold-changes**

| | | | | | RAW MEASUREMENT (mm) | | | | | | | | | | FOLD-CHANGE (from modern geoMean (mm)) | | | | | | | | | | |
|---|---|---|---|---|---|---|---|---|---|---|---|---|---|---|---|---|---|---|---|---|---|---|---|---|---|
| *SIDE* | *ELEMENT* | *AGE* | *COLLECTION* | *SAMPLE ID* | *MFL* | *MFD* | *FMD* | *DSA* | *FHH* | *FHW* | *FHSA* | *FBE* | *DML1* | *DML2* | *MFL (177.26)* | *MFD (8.73)* | *FMD (9.21)* | *DSA (623.65)* | *FHH (12.37)* | *FHW (12.22)* | *FHSA (347.59)* | *FBE (19.84)* | *DML1 (4.99)* | *DML2 (6.86)* | *FOLD-CHANGE (SIDED) AVERAGE* |
| Right | Femur | Modern | BMOC | 550 | 169.34 | 8.45 | 8.57 | 630.44 | 11.85 | 11.74 | 323.02 | 18.85 | 4.9 | 6.45 | -0.04 | -0.03 | -0.07 | 0.01 | -0.04 | -0.04 | -0.07 | -0.05 | -0.02 | -0.06 | -0.04 |
| Right | Femur | Modern | BMOC | 570 | 172.58 | 8.69 | 9.3 | 622.63 | 11.65 | 11.73 | 300.15 | 18.92 | 4.12 | 5.83 | -0.03 | 0 | 0.01 | 0 | -0.06 | -0.04 | -0.14 | -0.05 | -0.17 | -0.15 | -0.06 |
| Right | Femur | Modern | BMOC | BMOC001 | 187.31 | 8.22 | 9.31 | 633.68 | 12.21 | 11.86 | 328.32 | 20.16 | 5.09 | 6.77 | 0.06 | -0.06 | 0.01 | 0.02 | -0.01 | -0.03 | -0.06 | 0.02 | 0.02 | -0.01 | 0 |
| Right | Femur | Modern | BMOC | BMOC004 | 174.8 | 8.76 | 9.25 | 532.47 | 11.68 | 11.45 | 323.67 | 19.71 | 4.73 | 6.06 | -0.01 | 0 | 0 | -0.15 | -0.06 | -0.06 | -0.07 | -0.01 | -0.05 | -0.12 | -0.05 |
| Right | Femur | Modern | BMOC | BMOC005 | 186.45 | 9.14 | 9.81 | 593.86 | 12.88 | 12.47 | 374.67 | 20.32 | 4.87 | 7.2 | 0.05 | 0.05 | 0.06 | -0.05 | 0.04 | 0.02 | 0.08 | 0.02 | -0.02 | 0.05 | 0.03 |
| Right | Femur | Modern | BMOC | BMOC008 | NA | 8.55 | 9.49 | 646.02 | 13.64 | 13.4 | 425.78 | 20.53 | 5.4 | 6.43 | NA | -0.02 | 0.03 | 0.04 | 0.1 | 0.1 | 0.22 | 0.03 | 0.08 | -0.06 | 0.06 |
| Right | Femur | Modern | BMOC | BMOC014 | 171.89 | 8.67 | 9.22 | 650.34 | 12.07 | 12.26 | 323.44 | 20.53 | 5.4 | 6.97 | -0.03 | -0.01 | 0 | 0.04 | -0.02 | 0 | -0.07 | 0.03 | 0.08 | 0.02 | 0.01 |
| Right | Femur | Modern | BMOC | BMOC015 | 165.44 | 8.28 | 8.83 | 751.98 | 12.8 | 12.19 | 375.79 | 18.71 | 4.97 | 6.87 | -0.07 | -0.05 | -0.04 | 0.21 | 0.04 | 0 | 0.08 | -0.06 | 0 | 0 | 0.01 |
| Right | Femur | Modern | BMOC | BMOC019 | 180.28 | 9.07 | 9.43 | 744.49 | 12.57 | 12.33 | 360.5 | 20.91 | 5.38 | 7.35 | 0.02 | 0.04 | 0.02 | 0.19 | 0.02 | 0.01 | 0.04 | 0.05 | 0.08 | 0.07 | 0.05 |
| Right | Femur | Modern | BMOC | BMOC020 | 167.2 | 8.72 | 9.35 | 638.83 | 11.93 | 11.69 | 318.53 | 19.65 | 5.17 | 7.09 | -0.06 | 0 | 0.02 | 0.02 | -0.04 | -0.04 | -0.08 | -0.01 | 0.04 | 0.03 | -0.01 |
| Right | Femur | Modern | BMOC | BMOC021 | 175.3 | 8.89 | 9.02 | 532.14 | 12.26 | 12.01 | 338.87 | 19.5 | 4.75 | 7.1 | -0.01 | 0.02 | -0.02 | -0.15 | -0.01 | -0.02 | -0.03 | -0.02 | -0.05 | 0.03 | -0.02 |
| Right | Femur | Modern | BMOC | BMOC028 | NA | NA | NA | 665.23 | NA | NA | NA | 19.52 | 6.05 | 8.02 | NA | NA | NA | 0.07 | NA | NA | NA | -0.02 | 0.21 | 0.17 | 0.11 |
| Right | Femur | Modern | BMOC | BMOC030 | NA | 9.52 | 9.46 | NA | 12.72 | 12.43 | 363.45 | NA | NA | NA | NA | 0.09 | 0.03 | NA | 0.03 | 0.02 | 0.05 | NA | NA | NA | 0.04 |
| Right | Femur | Modern | BMOC | BMOC035 | 177.86 | 9.37 | 10.1 | 596.6 | 11.8 | 11.9 | 331.5 | 19.6 | 5.41 | 7.2 | 0 | 0.07 | 0.1 | -0.04 | -0.05 | -0.03 | -0.05 | -0.01 | 0.08 | 0.05 | 0.01 |
| Right | Femur | Modern | BMOC | BMOC073 | 174.02 | 8.37 | 9.4 | 584.62 | 12.72 | 12.67 | 359.59 | 19.54 | 5.17 | 6.96 | -0.02 | -0.04 | 0.02 | -0.06 | 0.03 | 0.04 | 0.03 | -0.01 | 0.04 | 0.01 | 0 |
| Right | Femur | Modern | BMOC | BMOC075 | NA | 8.1 | 8.28 | 643.69 | 11.9 | 12.1 | 354.95 | 19.13 | 4.48 | 6.5 | NA | -0.07 | -0.1 | 0.03 | -0.04 | -0.01 | 0.02 | -0.04 | -0.1 | -0.05 | -0.04 |
| Right | Femur | Modern | BMOC | BMOC137 | 186.91 | 8.93 | 9.99 | 641.76 | 13.53 | 13.4 | 408.93 | 21.86 | 5.29 | 8.03 | 0.05 | 0.02 | 0.08 | 0.03 | 0.09 | 0.1 | 0.18 | 0.1 | 0.06 | 0.17 | 0.09 |
| Right | Femur | Modern | BMOC | BMOC142 | 177.54 | 8.83 | 9.18 | 673.35 | 13.26 | 12.66 | 390.47 | 20.13 | 5.03 | 7.12 | 0 | 0.01 | 0 | 0.08 | 0.07 | 0.04 | 0.12 | 0.01 | 0.01 | 0.04 | 0.04 |
| Right | Femur | Modern | BMOC | BMOC156 | 180.38 | 10.02 | 9.14 | 564.04 | 11.97 | 11.86 | 314.01 | 19.85 | 4.55 | 6.76 | 0.02 | 0.15 | -0.01 | -0.1 | -0.03 | -0.03 | -0.1 | 0 | -0.09 | -0.02 | -0.02 |
| Right | Femur | Modern | BMOC | BMOC157 | 187.93 | 8.56 | 9.46 | 614.2 | 12.69 | 12.61 | 388.85 | 19.65 | 4.35 | 6.48 | 0.06 | -0.02 | 0.03 | -0.02 | 0.03 | 0.03 | 0.12 | -0.01 | -0.13 | -0.06 | 0 |
| Right | Femur | Modern | BMOC | BMOC169 | 179.64 | 8.76 | 9.22 | 639.66 | 12.87 | 13.15 | 405.4 | 21.07 | 5.16 | 6.95 | 0.01 | 0 | 0 | 0.03 | 0.04 | 0.08 | 0.17 | 0.06 | 0.03 | 0.01 | 0.04 |
| Right | Femur | Modern | BMOC | BMOC172 | 168.54 | 8.25 | 8.52 | 612.65 | 12.25 | 11.92 | 339.71 | 18.74 | 4.81 | 6.55 | -0.05 | -0.05 | -0.08 | -0.02 | -0.01 | -0.02 | -0.02 | -0.06 | -0.04 | -0.05 | -0.04 |
| Right | Femur | Modern | BMOC | BMOC173 | 181.07 | 8.12 | 8.8 | 577.39 | 12.05 | 12.23 | 310.19 | 19.87 | 4.93 | 6.91 | 0.02 | -0.07 | -0.04 | -0.07 | -0.03 | 0 | -0.11 | 0 | -0.01 | 0.01 | -0.03 |
| Right | Femur | Modern | BMOC | BMOC174 | 183.22 | 8.68 | 8.97 | 603.95 | 12.17 | 12.04 | 327.55 | 19.84 | 5.07 | 6.68 | 0.03 | -0.01 | -0.03 | -0.03 | -0.02 | -0.01 | -0.06 | 0 | 0.02 | -0.03 | -0.01 |
| Right | Femur | Modern | BMOC | BMOC197 | NA | NA | NA | NA | 11.6 | 11.37 | 296.7 | NA | NA | NA | NA | NA | NA | NA | -0.06 | -0.07 | -0.15 | NA | NA | NA | -0.09 |
| Right | Femur | Subfossil | TAO | TAO-66-21 | NA | NA | NA | NA | 13.62 | 13.31 | 428.74 | NA | NA | NA | NA | NA | NA | NA | 0.1 | 0.09 | 0.23 | NA | NA | NA | 0.14 |
| Right | Femur | Subfossil | TAO | TAO-66-25 | NA | NA | NA | NA | 12.9 | 13.01 | 352.67 | NA | NA | NA | NA | NA | NA | NA | 0.04 | 0.06 | 0.01 | NA | NA | NA | 0.04 |
| Right | Femur | Subfossil | TAO | TAO-66-26 | NA | NA | NA | NA | NA | NA | NA | NA | NA | 7 | NA | NA | NA | NA | NA | NA | NA | NA | NA | 0.02 | 0.02 |
| Right | Femur | Subfossil | TAO | TAO-66-29 | NA | NA | NA | NA | NA | NA | NA | 19.16 | NA | 7.38 | NA | NA | NA | NA | NA | NA | NA | -0.03 | NA | 0.08 | 0.02 |
| Right | Femur | Subfossil | TAO | TAO-66-32 | NA | NA | NA | NA | NA | NA | NA | 21.24 | NA | 8.12 | NA | NA | NA | NA | NA | NA | NA | 0.07 | NA | 0.18 | 0.13 |
| Right | Femur | Subfossil | TAO | TAO-66-33 | NA | NA | NA | NA | NA | NA | NA | NA | NA | 8.54 | NA | NA | NA | NA | NA | NA | NA | NA | NA | 0.24 | 0.24 |

| | | | | | RAW MEASUREMENT (mm) | | | | | | | | | | FOLD-CHANGE (from modern geoMean (mm)) | | | | | | | | | | |
|---|---|---|---|---|---|---|---|---|---|---|---|---|---|---|---|---|---|---|---|---|---|---|---|---|---|
| *SIDE* | *ELEMENT* | *AGE* | *COLLECTION* | *SAMPLE ID* | *MFL* | *MFD* | *FMD* | *DSA* | *FHH* | *FHW* | *FHSA* | *FBE* | *DML1* | *DML2* | *MFL (176.83)* | *MFD (8.55)* | *FMD (9.29)* | *DSA (565.14)* | *FHH (12.41)* | *FHW (12.36)* | *FHSA (347.28)* | *FBE (19.77)* | *DML1 (4.52)* | *DML2 (6.76)* | *FOLD-CHANGE (SIDED) AVERAGE* |
| Left | Femur | Modern | BMOC | 550 | 167.06 | 8.36 | 9.01 | 498.82 | 11.64 | 11.74 | 296.91 | 18.18 | 4.16 | 6.34 | -0.055 | -0.022 | -0.03 | -0.117 | -0.062 | -0.05 | -0.145 | -0.081 | -0.08 | -0.062 | -0.07 |
| Left | Femur | Modern | BMOC | 570 | 174.19 | 8.98 | 9.24 | 496.7 | 11.88 | 11.62 | 292.88 | 18.95 | 4.06 | 5.85 | -0.015 | 0.05 | -0.006 | -0.121 | -0.043 | -0.06 | -0.157 | -0.042 | -0.102 | -0.134 | -0.06 |
| Left | Femur | Modern | BMOC | BMOC001 | 187.48 | 8.12 | 9.41 | 588.34 | 12.2 | 12.22 | 336.07 | 19.62 | 5.24 | 6.25 | 0.06 | -0.05 | 0.013 | 0.041 | -0.017 | -0.011 | -0.032 | -0.008 | 0.159 | -0.075 | 0.01 |
| Left | Femur | Modern | BMOC | BMOC004 | 174.07 | 9.3 | 8.24 | 586.61 | 11.58 | 11.69 | 304.75 | 19.6 | 4.14 | 6.22 | -0.016 | 0.088 | -0.113 | 0.038 | -0.067 | -0.054 | -0.122 | -0.009 | -0.084 | -0.079 | -0.04 |
| Left | Femur | Modern | BMOC | BMOC005 | NA | 9.18 | 9.76 | 561.73 | NA | NA | NA | 20.31 | 4.05 | 7.36 | NA | 0.074 | 0.05 | -0.006 | NA | NA | NA | 0.027 | -0.104 | 0.089 | 0.02 |
| Left | Femur | Modern | BMOC | BMOC008 | 187.32 | 7.81 | 9.32 | 625.63 | 13.27 | 13.43 | 414.4 | 20.67 | 4.85 | 7.01 | 0.059 | -0.086 | 0.003 | 0.107 | 0.069 | 0.087 | 0.193 | 0.045 | 0.073 | 0.038 | 0.06 |
| Left | Femur | Modern | BMOC | BMOC014 | 172.19 | 8.1 | 9.6 | 547.06 | NA | NA | NA | 20.34 | 4.69 | 6.6 | -0.026 | -0.053 | 0.033 | -0.032 | NA | NA | NA | 0.029 | 0.037 | -0.023 | 0 |
| Left | Femur | Modern | BMOC | BMOC015 | 165.79 | 8.23 | 9.04 | 514.23 | 12.74 | 12.48 | 368.81 | 18.93 | 4.68 | 6.7 | -0.062 | -0.037 | -0.027 | -0.09 | 0.027 | 0.01 | 0.062 | -0.043 | 0.035 | -0.008 | -0.01 |
| Left | Femur | Modern | BMOC | BMOC019 | NA | 8.68 | 9.24 | NA | NA | NA | NA | NA | NA | NA | NA | 0.015 | -0.006 | NA | NA | NA | NA | NA | NA | NA | 0 |
| Left | Femur | Modern | BMOC | BMOC020 | 167.11 | 8.81 | 9.25 | 594.97 | 11.93 | 11.87 | 327.92 | 19.97 | 4.89 | 7.49 | -0.055 | 0.03 | -0.005 | 0.053 | -0.039 | -0.039 | -0.056 | 0.01 | 0.082 | 0.109 | 0.01 |
| Left | Femur | Modern | BMOC | BMOC021 | 175.14 | 8.58 | 9.2 | 540.57 | 12.14 | 12.23 | 351.3 | 19.64 | 4.31 | 6.99 | -0.01 | 0.004 | -0.01 | -0.043 | -0.022 | -0.01 | 0.012 | -0.007 | -0.047 | 0.035 | -0.01 |
| Left | Femur | Modern | BMOC | BMOC028 | 185.84 | 8.27 | 10.58 | 507.32 | 12.63 | 12.4 | 350.76 | 19.85 | 5.18 | 7.44 | 0.051 | -0.033 | 0.138 | -0.102 | 0.018 | 0.003 | 0.01 | 0.004 | 0.146 | 0.101 | 0.03 |
| Left | Femur | Modern | BMOC | BMOC073 | 174.51 | 8.57 | 9.19 | 654.95 | 12.3 | 12.44 | 344.67 | 19.31 | 4.65 | 6.62 | -0.013 | 0.002 | -0.011 | 0.159 | -0.009 | 0.007 | -0.008 | -0.023 | 0.028 | -0.02 | 0.01 |
| Left | Femur | Modern | BMOC | BMOC075 | 171.84 | 8.25 | 8.34 | 572.46 | 11.88 | 12.26 | 350.31 | 19.64 | 4.36 | 6.27 | -0.028 | -0.035 | -0.103 | 0.013 | -0.043 | -0.008 | 0.009 | -0.007 | -0.036 | -0.072 | -0.03 |
| Left | Femur | Modern | BMOC | BMOC137 | 188.25 | 9.58 | 10.09 | 688.31 | 13.67 | 13.57 | 419.48 | 21.66 | 4.84 | 7.61 | 0.065 | 0.121 | 0.086 | 0.218 | 0.102 | 0.098 | 0.208 | 0.096 | 0.07 | 0.126 | 0.12 |
| Left | Femur | Modern | BMOC | BMOC142 | 177.38 | 8.33 | 9.27 | 519.6 | 13.22 | 12.92 | 380.2 | 20.14 | 4.35 | 7.62 | 0.003 | -0.026 | -0.003 | -0.081 | 0.065 | 0.046 | 0.095 | 0.019 | -0.038 | 0.128 | 0.02 |
| Left | Femur | Modern | BMOC | BMOC157 | 189.02 | 8.51 | 9.83 | 623.03 | 12.69 | 12.67 | 363.94 | 20.1 | 4.3 | 6.43 | 0.069 | -0.005 | 0.058 | 0.102 | 0.023 | 0.025 | 0.048 | 0.017 | -0.049 | -0.048 | 0.02 |
| Left | Femur | Modern | BMOC | BMOC163 | NA | 9.32 | 11.34 | NA | 13.45 | 13.48 | 409.19 | NA | NA | NA | NA | 0.09 | 0.22 | NA | 0.084 | 0.091 | 0.178 | NA | NA | NA | 0.13 |
| Left | Femur | Modern | BMOC | BMOC172 | 168.57 | 8.35 | 8.26 | 524 | 12.01 | 11.76 | 312.55 | 18.9 | 4.26 | 6.64 | -0.047 | -0.023 | -0.111 | -0.073 | -0.032 | -0.048 | -0.1 | -0.044 | -0.058 | -0.017 | -0.06 |
| Left | Femur | Modern | BMOC | BMOC173 | 181.08 | 8.14 | 8.67 | 523.1 | 12.26 | 12.2 | 327.34 | 19.9 | 4.62 | 7.03 | 0.024 | -0.048 | -0.067 | -0.074 | -0.012 | -0.013 | -0.057 | 0.006 | 0.022 | 0.041 | -0.02 |
| Left | Femur | Modern | BMOC | BMOC174 | 183.46 | 8.72 | 9.25 | 521.47 | 12.32 | 12.24 | 346.58 | 19.72 | 4.61 | 6.93 | 0.038 | 0.02 | -0.005 | -0.077 | -0.007 | -0.009 | -0.002 | -0.003 | 0.02 | 0.026 | 0 |
| Left | Femur | Modern | BMOC | BMOC180 | NA | 8.52 | 9.71 | NA | 12.88 | 12.81 | 382.92 | NA | NA | NA | NA | -0.003 | 0.045 | NA | 0.038 | 0.037 | 0.103 | NA | NA | NA | 0.04 |
| Left | Femur | Modern | BMOC | BMOC191 | NA | 8.48 | 8.9 | 642.44 | NA | NA | NA | 20.53 | 4.71 | 6.67 | NA | -0.008 | -0.042 | 0.137 | NA | NA | NA | 0.038 | 0.042 | -0.013 | 0.03 |
| Left | Femur | Modern | BMOC | BMOC197 | 172.51 | 8.25 | 8.88 | 589.82 | 11.76 | 11.41 | 303.97 | 19.52 | 4.26 | 6.2 | -0.024 | -0.035 | -0.044 | 0.044 | -0.052 | -0.077 | -0.125 | -0.013 | -0.058 | -0.082 | -0.05 |
| Left | Femur | Subfossil | TAO | TAO-66-23 | NA | NA | NA | NA | 13.04 | 13.67 | 389.45 | NA | NA | NA | NA | NA | NA | NA | 0.051 | 0.106 | 0.121 | NA | NA | NA | 0.09 |
| Left | Femur | Subfossil | TAO | TAO-66-24 | NA | NA | NA | NA | 12.99 | 12.93 | 348.53 | NA | NA | NA | NA | NA | NA | NA | 0.047 | 0.046 | 0.004 | NA | NA | NA | 0.03 |
| Left | Femur | Subfossil | TAO | TAO-66-30 | NA | NA | NA | 658.3 | NA | NA | NA | 20.5 | 5.31 | 7.04 | NA | NA | NA | 0.165 | NA | NA | NA | 0.037 | 0.174 | 0.042 | 0.1 |
| Left | Femur | Subfossil | TAO | TAO-66-31 | NA | NA | NA | NA | NA | NA | NA | NA | NA | 5.59 | NA | NA | NA | NA | NA | NA | NA | NA | NA | -0.173 | -0.17 |
| Left | Femur | Subfossil | TAO | TAO-66-34 | NA | NA | NA | NA | NA | NA | NA | NA | 5.23 | 7.57 | NA | NA | NA | NA | NA | NA | NA | NA | 0.157 | 0.12 | 0.14 |

| | | | | | RAW MEASUREMENT (mm) | | | | | | | FOLD-CHANGE (from modern geoMean (mm)) | | | | | | | |
|---|---|---|---|---|---|---|---|---|---|---|---|---|---|---|---|---|---|---|---|
| *SIDE* | *ELEMENT* | *AGE* | *COLLECTION* | *SAMPLE ID* | *MHL* | *MHD1* | *MHD2* | *VHD* | *HHW* | *BBH* | *HHSA* | *MHL (91.23)* | *MHD1 (7.42)* | *1HD2 (6.12* | *VHD (11.29)* | *HHW (9.21)* | *BBH (20.92)* | *HHSA (173.15)* | *FOLD-CHANGE (SIDED) AVERAGE* |
| Right | Humerus | Modern | BMOC | 550 | 88.45 | 6.67 | 6.05 | 10.69 | 8.93 | 20.47 | 154.76 | -0.03 | -0.1 | -0.01 | -0.05 | -0.03 | -0.02 | -0.11 | -0.05 |
| Right | Humerus | Modern | BMOC | 570 | 89.62 | 7.7 | 5.98 | 10.69 | 8.53 | 20.01 | 144.88 | -0.02 | 0.04 | -0.02 | -0.05 | -0.07 | -0.04 | -0.16 | -0.05 |
| Right | Humerus | Modern | BMOC | BMOC001 | 95.06 | 6.87 | 5.95 | 11.12 | 9.34 | 21.24 | 168.95 | 0.04 | -0.07 | -0.03 | -0.02 | 0.01 | 0.02 | -0.02 | -0.01 |
| Right | Humerus | Modern | BMOC | BMOC004 | NA | 7.06 | 6.61 | 9.03 | 8.81 | 21.38 | 116.03 | NA | -0.05 | 0.08 | -0.2 | -0.04 | 0.02 | -0.33 | -0.09 |
| Right | Humerus | Modern | BMOC | BMOC008 | 94.86 | 7.9 | 6.56 | 12.63 | 10.29 | 21.02 | 225.5 | 0.04 | 0.06 | 0.07 | 0.12 | 0.12 | 0 | 0.3 | 0.1 |
| Right | Humerus | Modern | BMOC | BMOC015 | 86.69 | 8.31 | 6.5 | 11.92 | 9.6 | 21.12 | 192.26 | -0.05 | 0.12 | 0.06 | 0.06 | 0.04 | 0.01 | 0.11 | 0.05 |
| Right | Humerus | Modern | BMOC | BMOC019 | 92.8 | 7.87 | 6.22 | 11.75 | 9.36 | 21.78 | 188.04 | 0.02 | 0.06 | 0.02 | 0.04 | 0.02 | 0.04 | 0.09 | 0.04 |
| Right | Humerus | Modern | BMOC | BMOC020 | 88.96 | 7.03 | 5.81 | 11.52 | 9.55 | 20.99 | 176.23 | -0.02 | -0.05 | -0.05 | 0.02 | 0.04 | 0 | 0.02 | -0.01 |
| Right | Humerus | Modern | BMOC | BMOC028 | 92.47 | 6.68 | 5.32 | 12.01 | 9.11 | 20.64 | 178.42 | 0.01 | -0.1 | -0.13 | 0.06 | -0.01 | -0.01 | 0.03 | -0.02 |
| Right | Humerus | Modern | BMOC | BMOC030 | NA | 7.53 | 6.66 | 11.05 | 8.85 | NA | 183.25 | NA | 0.01 | 0.09 | -0.02 | -0.04 | NA | 0.06 | 0.02 |
| Right | Humerus | Modern | BMOC | BMOC043 | 89.43 | 6.26 | 5.63 | 11.25 | 8.89 | 20.75 | 163.59 | -0.02 | -0.16 | -0.08 | 0 | -0.03 | -0.01 | -0.06 | -0.05 |
| Right | Humerus | Modern | BMOC | BMOC073 | 89.18 | 8.39 | 6.06 | 12.05 | 9.2 | 21.27 | 192.9 | -0.02 | 0.13 | -0.01 | 0.07 | 0 | 0.02 | 0.11 | 0.04 |
| Right | Humerus | Modern | BMOC | BMOC075 | NA | 7.56 | 6.07 | NA | NA | 19.93 | NA | NA | 0.02 | -0.01 | NA | NA | -0.05 | NA | -0.01 |
| Right | Humerus | Modern | BMOC | BMOC125 | 98.47 | 7.39 | 5.87 | 11.76 | 10.15 | 22.29 | 200.736 | 0.08 | 0 | -0.04 | 0.04 | 0.1 | 0.07 | 0.16 | 0.06 |
| Right | Humerus | Modern | BMOC | BMOC137 | 96.96 | 7.8 | 6.55 | 12.55 | 10.37 | 23.85 | 206.07 | 0.06 | 0.05 | 0.07 | 0.11 | 0.13 | 0.14 | 0.19 | 0.11 |
| Right | Humerus | Modern | BMOC | BMOC142 | 92.06 | 7.8 | 5.86 | 11.91 | 9.37 | 21.52 | 181.74 | 0.01 | 0.05 | -0.04 | 0.05 | 0.02 | 0.03 | 0.05 | 0.02 |
| Right | Humerus | Modern | BMOC | BMOC157 | NA | 7.4 | 6.46 | NA | NA | 20.36 | NA | NA | 0 | 0.06 | NA | NA | -0.03 | NA | 0.01 |
| Right | Humerus | Modern | BMOC | BMOC172 | 85.54 | 7.55 | 5.76 | 10.85 | 9.3 | 19.77 | 174.64 | -0.06 | 0.02 | -0.06 | -0.04 | 0.01 | -0.06 | 0.01 | -0.03 |
| Right | Humerus | Modern | BMOC | BMOC173 | 91.08 | 7.31 | 6.08 | 10.91 | 8.58 | 20.67 | 152.42 | 0 | -0.01 | -0.01 | -0.03 | -0.07 | -0.01 | -0.12 | -0.04 |
| Right | Humerus | Modern | BMOC | BMOC174 | 92.6 | 7.01 | 6.08 | 11.31 | 8.93 | 20.8 | 176.8 | 0.02 | -0.06 | -0.01 | 0 | -0.03 | -0.01 | 0.02 | -0.01 |
| Right | Humerus | Modern | BMOC | BMOC180 | NA | 7.7 | 6.21 | NA | NA | 21.09 | NA | NA | 0.04 | 0.02 | NA | NA | 0.01 | NA | 0.02 |
| Right | Humerus | Modern | BMOC | BMOC197 | 89.99 | 7.37 | 6.22 | 10.47 | 8.44 | 20.19 | 143.86 | -0.01 | -0.01 | 0.02 | -0.07 | -0.08 | -0.03 | -0.17 | -0.05 |
| Right | Humerus | Modern | BMOC | Fogel 2012 | 88.98 | 7.92 | 6.37 | 10.97 | 8.85 | 19.56 | 176.91 | -0.02 | 0.07 | 0.04 | -0.03 | -0.04 | -0.07 | 0.02 | 0 |
| Right | Humerus | Subfossil | TAO | TAO-66-2 | NA | NA | NA | 14.21 | 10.61 | NA | 248.06 | NA | NA | NA | 0.26 | 0.15 | NA | 0.43 | 0.28 |

| | | | | | RAW MEASUREMENT (mm) | | | | | | | FOLD-CHANGE (from modern geoMean (mm)) | | | | | | | |
|---|---|---|---|---|---|---|---|---|---|---|---|---|---|---|---|---|---|---|---|
| *SIDE* | *ELEMENT* | *AGE* | *COLLECTION* | *SAMPLE ID* | *MHL* | *MHD1* | *MHD2* | *VHD* | *HHW* | *BBH* | *HHSA* | *MHL (90.79)* | *MHD1 (7.44)* | *1HD2 (6.11* | *VHD (11.41)* | *HHW (9.19)* | *BBH (20.90)* | *HHSA (177.67)* | *FOLD-CHANGE (SIDED) AVERAGE* |
| Left | Humerus | Modern | BMOC | 550 | 88.15 | 6.69 | 5.54 | 10.71 | 8.28 | 20.22 | 153.45 | -0.03 | -0.1 | -0.09 | -0.06 | -0.1 | -0.03 | -0.14 | -0.08 |
| Left | Humerus | Modern | BMOC | 570 | 90.23 | 7.65 | 5.99 | 10.5 | 8.55 | 19.74 | 157.34 | -0.01 | 0.03 | -0.02 | -0.08 | -0.07 | -0.06 | -0.11 | -0.05 |
| Left | Humerus | Modern | BMOC | BMOC001 | 94.98 | 7.06 | 5.92 | 11.78 | 9.51 | 21.28 | 190.85 | 0.05 | -0.05 | -0.03 | 0.03 | 0.03 | 0.02 | 0.07 | 0.02 |
| Left | Humerus | Modern | BMOC | BMOC004 | 90.7 | 6.87 | 5.91 | 10.84 | 8.68 | 21.53 | 157.92 | 0 | -0.08 | -0.03 | -0.05 | -0.06 | 0.03 | -0.11 | -0.04 |
| Left | Humerus | Modern | BMOC | BMOC008 | 96.63 | 7.7 | 6.57 | 12.23 | 9.89 | 21.4 | 203.5 | 0.06 | 0.03 | 0.07 | 0.07 | 0.08 | 0.02 | 0.15 | 0.07 |
| Left | Humerus | Modern | BMOC | BMOC015 | 86.62 | 8.8 | 6.32 | 11.82 | 9.89 | 21.46 | 198.75 | -0.05 | 0.18 | 0.03 | 0.04 | 0.08 | 0.03 | 0.12 | 0.06 |
| Left | Humerus | Modern | BMOC | BMOC019 | 92.74 | 7.53 | 6.35 | 12.25 | 10 | 21.36 | 193.16 | 0.02 | 0.01 | 0.04 | 0.07 | 0.09 | 0.02 | 0.09 | 0.05 |
| Left | Humerus | Modern | BMOC | BMOC020 | 88.67 | 7.16 | 5.92 | 11.54 | 9.68 | 21.4 | 176.89 | -0.02 | -0.04 | -0.03 | 0.01 | 0.05 | 0.02 | 0 | 0 |
| Left | Humerus | Modern | BMOC | BMOC028 | 91.96 | 6.77 | 5.22 | 11.72 | 9.22 | 20.91 | 175.46 | 0.01 | -0.09 | -0.15 | 0.03 | 0 | 0 | -0.01 | -0.03 |
| Left | Humerus | Modern | BMOC | BMOC043 | NA | NA | NA | NA | NA | 20.72 | NA | NA | NA | NA | NA | NA | -0.01 | NA | -0.01 |
| Left | Humerus | Modern | BMOC | BMOC073 | 89.01 | 7.95 | 6.15 | 11.72 | 9.17 | 21.19 | 181.07 | -0.02 | 0.07 | 0.01 | 0.03 | 0 | 0.01 | 0.02 | 0.02 |
| Left | Humerus | Modern | BMOC | BMOC075 | 87.49 | 7.41 | 6 | 11.06 | 8.83 | 19.92 | 173.39 | -0.04 | 0 | -0.02 | -0.03 | -0.04 | -0.05 | -0.02 | -0.03 |
| Left | Humerus | Modern | BMOC | BMOC137 | 96.75 | 7.69 | 6.58 | 12.56 | 10.4 | 23.62 | 205.02 | 0.07 | 0.03 | 0.08 | 0.1 | 0.13 | 0.13 | 0.15 | 0.1 |
| Left | Humerus | Modern | BMOC | BMOC142 | 92.11 | 7.96 | 5.98 | 11.93 | 9.33 | 21.33 | 190.92 | 0.01 | 0.07 | -0.02 | 0.05 | 0.01 | 0.02 | 0.07 | 0.03 |
| Left | Humerus | Modern | BMOC | BMOC157 | NA | 7.45 | 6.31 | 11.12 | 8.51 | NA | 172.16 | NA | 0 | 0.03 | -0.03 | -0.07 | NA | -0.03 | -0.02 |
| Left | Humerus | Modern | BMOC | BMOC172 | 85.93 | 7.49 | 5.52 | 11.38 | 9.26 | 19.56 | 185.17 | -0.05 | 0.01 | -0.1 | 0 | 0.01 | -0.06 | 0.04 | -0.02 |
| Left | Humerus | Modern | BMOC | BMOC173 | 90.96 | 7.48 | 6.09 | 11.29 | 8.97 | 20.66 | 165.49 | 0 | 0.01 | 0 | -0.01 | -0.02 | -0.01 | -0.07 | -0.02 |
| Left | Humerus | Modern | BMOC | BMOC174 | 89.67 | 7.84 | 6.47 | 11.54 | 8.77 | 20.99 | 173.95 | -0.01 | 0.05 | 0.06 | 0.01 | -0.05 | 0 | -0.02 | 0.01 |
| Left | Humerus | Modern | BMOC | BMOC180 | 92.92 | 6.75 | 7.37 | 11.21 | 9.85 | 20.96 | 196.03 | 0.02 | -0.09 | 0.21 | -0.02 | 0.07 | 0 | 0.1 | 0.04 |
| Left | Humerus | Modern | BMOC | BMOC197 | 90.74 | 7.52 | 6.33 | 10.89 | 8.72 | 20.34 | 150.57 | 0 | 0.01 | 0.04 | -0.05 | -0.05 | -0.03 | -0.15 | -0.03 |
| Left | Humerus | Modern | BMOC | Fogel 2012 | 89.63 | 7.37 | 6.02 | 10.44 | 8.71 | 19.7 | 167.27 | -0.01 | -0.01 | -0.02 | -0.09 | -0.05 | -0.06 | -0.06 | -0.04 |

**Supplementary Table 4: Average percent differences between radial caliper and 3D model measurements**

| | Modern | | Subfossil | | % Difference |
|---|---|---|---|---|---|
| | **% Difference** | **Standard Deviation** | **% Difference** | **Standard Deviation** | **t-test p** |
| **Right Femur** | -0.03 | 3.02 | 1.66 | 0.58 | 0.0465 |
| **Left Femur** | 0.8 | 1.74 | 1.54 | 0.66 | 0.5851 |
| **Right Humerus** | -2.02 | 7.04 | 0.41 | 1.67 | N/A |
| **Left Humerus** | -1.48 | 6.47 | N/A | N/A | N/A |

**Supplementary Table 5: Radial caliper versus 3D scan Avizo measurements**

| | | | | | *RAW MEASUREMENT % DIFFERENCE (mm)* | | | | | | |
|---|---|---|---|---|---|---|---|---|---|---|---|
| *SIDE* | *ELEMENT* | *AGE* | *COLLECTION* | *SAMPLE ID* | *MFL* | *MFD* | *FHH* | *FHW* | *FBE* | *AVE %* | *STDEV* |
| Right | Femur | Modern | BMOC | 550 | 0 | 1.64 | -1.19 | 2.19 | 0.58 | 0.65 | 1.34 |
| Right | Femur | Modern | BMOC | 570 | 0.08 | 4.06 | 0.09 | 0.93 | 2.81 | 1.59 | 1.77 |
| Right | Femur | Modern | BMOC | BMOC001 | 0.05 | 0.24 | -2.4 | 2.83 | 0.59 | 0.26 | 1.86 |
| Right | Femur | Modern | BMOC | BMOC004 | -0.52 | -2.9 | -1.55 | 3.18 | 1.76 | 0 | 2.46 |
| Right | Femur | Modern | BMOC | BMOC005 | -0.54 | -0.11 | -2.75 | 3.62 | 1.08 | 0.26 | 2.34 |
| Right | Femur | Modern | BMOC | BMOC008 | | 2.08 | -3.35 | 2.51 | 0.87 | 0.53 | 2.68 |
| Right | Femur | Modern | BMOC | BMOC014 | 0.17 | -0.58 | -1.08 | 1.38 | 0.05 | -0.01 | 0.93 |
| Right | Femur | Modern | BMOC | BMOC015 | 0.63 | -0.73 | -0.78 | 2.99 | 1.96 | 0.81 | 1.66 |
| Right | Femur | Modern | BMOC | BMOC019 | 0.25 | -2 | 0.24 | -0.49 | 1.52 | -0.1 | 1.29 |
| Right | Femur | Modern | BMOC | BMOC020 | -3.23 | 0 | -1.01 | 3.2 | -0.2 | -0.25 | 2.31 |
| Right | Femur | Modern | BMOC | BMOC021 | 10.8 | -1.59 | -1.07 | 2.14 | 1.07 | 2.27 | 5.01 |
| Right | Femur | Modern | BMOC | BMOC028 | | | | | 2.73 | 2.73 | NA |
| Right | Femur | Modern | BMOC | BMOC030 | | -2.77 | -3.03 | 3.09 | | -0.9 | 3.46 |
| Right | Femur | Modern | BMOC | BMOC035 | 0.3 | -4.47 | 0.08 | 0.75 | -0.77 | -0.82 | 2.12 |
| Right | Femur | Modern | BMOC | BMOC073 | 0.08 | 1.78 | -0.63 | 0.32 | 1.88 | 0.68 | 1.1 |
| Right | Femur | Modern | BMOC | BMOC075 | | 2.8 | 0.42 | 2.85 | 4.6 | 2.67 | 1.72 |
| Right | Femur | Modern | BMOC | BMOC137 | -0.18 | 1.22 | -2.39 | 1.85 | 0.32 | 0.16 | 1.63 |
| Right | Femur | Modern | BMOC | BMOC142 | -0.34 | 2.46 | -0.53 | 4.93 | -0.35 | 1.24 | 2.41 |
| Right | Femur | Modern | BMOC | BMOC156 | 0.14 | -1.2 | 1.25 | -1.1 | 2.39 | 0.29 | 1.54 |
| Right | Femur | Modern | BMOC | BMOC157 | 0.23 | 3.78 | -2.39 | 2.35 | 0.96 | 0.99 | 2.33 |
| Right | Femur | Modern | BMOC | BMOC169 | -0.21 | 1.47 | -46.08 | -45.68 | 2.02 | -17.69 | 25.74 |
| Right | Femur | Modern | BMOC | BMOC172 | 0.27 | 0.84 | -3.07 | 0.67 | 2.74 | 0.29 | 2.1 |
| Right | Femur | Modern | BMOC | BMOC173 | 0.16 | 2.67 | -0.17 | 0.41 | -0.15 | 0.58 | 1.19 |
| Right | Femur | Modern | BMOC | BMOC174 | -0.28 | 2.05 | -0.41 | 1.81 | 4.05 | 1.44 | 1.85 |
| Right | Femur | Modern | BMOC | BMOC197 | | | 0.43 | 2.69 | | 1.56 | 1.6 |
| Right | Femur | Subfossil | TAO | TAO-66-21 | | | 1.75 | 2.3 | | 2.02 | 0.39 |
| Right | Femur | Subfossil | TAO | TAO-66-25 | | | 2.37 | 1.3 | | 1.84 | 0.76 |
| Right | Femur | Subfossil | TAO | TAO-66-32 | | | | | 1.12 | 1.12 | NA |

| | | | | | *RAW MEASUREMENT % DIFFERENCE (mm)* | | | | | | |
|---|---|---|---|---|---|---|---|---|---|---|---|
| *SIDE* | *ELEMENT* | *AGE* | *COLLECTION* | *SAMPLE ID* | *MFL* | *MFD* | *FHH* | *FHW* | *FBE* | *AVE %* | *STDEV* |
| Left | Femur | Modern | BMOC | 550 | 0.17 | -1.45 | 1.62 | 2.52 | 6.54 | 1.88 | 3.01 |
| Left | Femur | Modern | BMOC | 570 | -0.01 | 0.67 | -1.19 | 2.72 | 2.71 | 0.98 | 1.71 |
| Left | Femur | Modern | BMOC | BMOC001 | 0.09 | -0.87 | -0.49 | 0.49 | 2.02 | 0.25 | 1.12 |
| Left | Femur | Modern | BMOC | BMOC004 | 0.13 | -4.4 | -0.61 | 1.19 | 2.62 | -0.21 | 2.63 |
| Left | Femur | Modern | BMOC | BMOC005 | | -0.88 | | | 0.54 | -0.17 | 1 |
| Left | Femur | Modern | BMOC | BMOC008 | 0.17 | 8.46 | 0.9 | 1.04 | -0.15 | 2.08 | 3.6 |
| Left | Femur | Modern | BMOC | BMOC014 | | 1.47 | | | 1.85 | 1.66 | 0.27 |
| Left | Femur | Modern | BMOC | BMOC015 | 0.08 | 2.04 | 0.47 | 1.9 | 0.26 | 0.95 | 0.94 |
| Left | Femur | Modern | BMOC | BMOC019 | | 1.94 | | | | 1.94 | NA |
| Left | Femur | Modern | BMOC | BMOC020 | -0.04 | -1.26 | 0.33 | 2.91 | -0.65 | 0.26 | 1.6 |
| Left | Femur | Modern | BMOC | BMOC021 | -0.12 | 0.47 | 0.74 | 1.14 | -0.41 | 0.36 | 0.63 |
| Left | Femur | Modern | BMOC | BMOC028 | -0.02 | 3.21 | -0.87 | 2.39 | 2.54 | 1.45 | NA |
| Left | Femur | Modern | BMOC | BMOC073 | 0.08 | 0.58 | -0.82 | 0.88 | 2.3 | 0.61 | 1.15 |
| Left | Femur | Modern | BMOC | BMOC075 | -0.17 | 0.48 | 1.75 | 1.7 | 2.41 | 1.23 | 1.05 |
| Left | Femur | Modern | BMOC | BMOC137 | -0.63 | -3.18 | -1.92 | -0.07 | 0.46 | -1.07 | 1.48 |
| Left | Femur | Modern | BMOC | BMOC142 | -0.02 | 5.26 | -0.91 | 2.22 | 0.54 | 1.42 | 2.43 |
| Left | Femur | Modern | BMOC | BMOC157 | 0 | 1.4 | -2.31 | 1.33 | 3.09 | 0.7 | 2.01 |
| Left | Femur | Modern | BMOC | BMOC163 | | 1.07 | -0.6 | 1.33 | | 0.6 | 1.04 |
| Left | Femur | Modern | BMOC | BMOC172 | -0.02 | 2.37 | -0.5 | 1.02 | -0.11 | 0.55 | 1.16 |
| Left | Femur | Modern | BMOC | BMOC173 | -0.18 | 2.07 | -0.65 | 1.14 | 1.55 | 0.78 | 1.16 |
| Left | Femur | Modern | BMOC | BMOC174 | -0.16 | 2.04 | 0.24 | 1.22 | 4.41 | 1.55 | 1.82 |
| Left | Femur | Modern | BMOC | BMOC180 | | 5.92 | -6.91 | 0.08 | | -0.3 | 6.42 |
| Left | Femur | Modern | BMOC | BMOC191 | | 0.82 | | | 0.15 | 0.48 | 0.48 |
| Left | Femur | Modern | BMOC | BMOC197 | -0.05 | 3.1 | -0.51 | 2.42 | 1.07 | 1.21 | 1.55 |
| Left | Femur | Subfossil | TAO | TAO-66-23 | | | 0.15 | -1.1 | | -0.48 | 0.89 |
| Left | Femur | Subfossil | TAO | TAO-66-24 | | | 1.91 | 1.31 | | 1.61 | 0.42 |
| Left | Femur | Subfossil | TAO | TAO-66-30 | | | | | 3.5 | 3.5 | NA |
| Left | Femur | Subfossil | TAO | TAO-66-34 | | | | | | NA | NA |

| | | | | | *RAW MEASUREMENT % DIFFERENCE (mm)* | | | | | | |
|---|---|---|---|---|---|---|---|---|---|---|---|
| *SIDE* | *ELEMENT* | *AGE* | *COLLECTION* | *SAMPLE ID* | *MHL* | *MHD* | *VHD* | *HHW* | *BBH* | *AVE %* | *STDEV* |
| Right | Humerus | Modern | BMOC | 550 | 0.11 | -3.82 | 1.85 | 3.63 | -0.29 | 0.3 | 2.78 |
| Right | Humerus | Modern | BMOC | 570 | 0.04 | -21.9 | 1.12 | 5.36 | -1.26 | -3.33 | 10.68 |
| Right | Humerus | Modern | BMOC | BMOC001 | -0.56 | -8.34 | -1.54 | 3.06 | 0.98 | -1.28 | 4.31 |
| Right | Humerus | Modern | BMOC | BMOC004 | | -3.31 | 11.58 | 2.36 | -1.94 | 2.17 | 6.72 |
| Right | Humerus | Modern | BMOC | BMOC008 | -0.24 | -10.11 | -3.46 | 1.45 | | -3.09 | 5.1 |
| Right | Humerus | Modern | BMOC | BMOC015 | 0.09 | -14.59 | -3.67 | 5.37 | 0.9 | -2.38 | 7.54 |
| Right | Humerus | Modern | BMOC | BMOC019 | -0.13 | -9.45 | -3.02 | 8.98 | -1.34 | -0.99 | 6.63 |
| Right | Humerus | Modern | BMOC | BMOC020 | -0.11 | 0.57 | -7.66 | 2.48 | -0.38 | -1.02 | 3.88 |
| Right | Humerus | Modern | BMOC | BMOC028 | 0.02 | -18.84 | 0.42 | 3.03 | 1.01 | -2.87 | 9 |
| Right | Humerus | Modern | BMOC | BMOC030 | | 0.93 | -0.91 | 8.44 | | 2.82 | 4.95 |
| Right | Humerus | Modern | BMOC | BMOC043 | 0.12 | 12.99 | -5.29 | 5.47 | 1.1 | 2.88 | 6.83 |
| Right | Humerus | Modern | BMOC | BMOC073 | 0.15 | -24.03 | -7.76 | 4.57 | -0.38 | -5.49 | 11.27 |
| Right | Humerus | Modern | BMOC | BMOC075 | | -3.23 | | | 1.05 | -1.09 | 3.02 |
| Right | Humerus | Modern | BMOC | BMOC125 | -0.64 | -14.36 | -2.15 | 7.22 | 1.43 | -1.7 | 7.92 |
| Right | Humerus | Modern | BMOC | BMOC137 | 0.02 | -11.81 | -0.8 | 3.04 | -4.5 | -2.81 | 5.7 |
| Right | Humerus | Modern | BMOC | BMOC142 | -0.17 | -18.33 | 1.67 | 5.1 | 0 | -2.35 | 9.18 |
| Right | Humerus | Modern | BMOC | BMOC157 | | -14.34 | | | 0.64 | -6.85 | 10.59 |
| Right | Humerus | Modern | BMOC | BMOC172 | -0.25 | -19.32 | -12.33 | 4.11 | -0.41 | -5.64 | 9.79 |
| Right | Humerus | Modern | BMOC | BMOC173 | 0.21 | -14.06 | 3.16 | 4.89 | -0.05 | -1.17 | 7.49 |
| Right | Humerus | Modern | BMOC | BMOC174 | -0.08 | -8.79 | 2.1 | 1.22 | 1.48 | -0.81 | 4.53 |
| Right | Humerus | Modern | BMOC | BMOC180 | | -13.44 | | | -1.19 | -7.32 | 8.66 |
| Right | Humerus | Modern | BMOC | BMOC197 | 0.07 | -7.9 | 3.84 | 4.29 | 0.2 | 0.1 | 4.89 |
| Right | Humerus | Modern | BMOC | Fogel 2012 | -0.38 | -22.94 | 0.27 | 1.68 | -1.08 | -4.49 | 10.36 |
| Right | Humerus | Subfossil | TAO | TAO-66-2 | | | -0.78 | 1.59 | | 0.41 | 1.67 |

| | | | | | *RAW MEASUREMENT % DIFFERENCE (mm)* | | | | | | |
|---|---|---|---|---|---|---|---|---|---|---|---|
| *SIDE* | *ELEMENT* | *AGE* | *COLLECTION* | *SAMPLE ID* | *MHL* | *MHD* | *VHD* | *HHW* | *BBH* | *AVE %* | *STDEV* |
| Left | Humerus | Modern | BMOC | 550 | 0.08 | -3.03 | 4.2 | 7.67 | 1.86 | 2.16 | 4.06 |
| Left | Humerus | Modern | BMOC | 570 | -0.14 | -14.29 | 1.89 | 2.2 | 2.06 | -1.66 | 7.12 |
| Left | Humerus | Modern | BMOC | BMOC001 | -0.65 | -13.62 | -6.4 | 3.41 | 1.68 | -3.12 | 6.94 |
| Left | Humerus | Modern | BMOC | BMOC004 | 0.1 | -0.15 | -2.14 | 4.5 | -0.09 | 0.44 | 2.45 |
| Left | Humerus | Modern | BMOC | BMOC008 | -0.65 | -14.19 | 1.3 | 5.51 | -1.89 | -1.98 | 7.38 |
| Left | Humerus | Modern | BMOC | BMOC015 | -0.07 | -14.5 | -2.14 | 0.81 | -1.64 | -3.51 | 6.26 |
| Left | Humerus | Modern | BMOC | BMOC019 | -0.42 | -2.15 | -7.01 | 2.47 | -0.61 | -1.54 | 3.48 |
| Left | Humerus | Modern | BMOC | BMOC020 | 0.05 | -4.43 | -5.98 | 2.15 | -4 | -2.44 | 3.4 |
| Left | Humerus | Modern | BMOC | BMOC028 | 0.02 | -22.33 | 3.03 | 3.73 | -0.77 | -3.27 | 10.83 |
| Left | Humerus | Modern | BMOC | BMOC043 | | | | | 0.77 | 0.77 | NA |
| Left | Humerus | Modern | BMOC | BMOC073 | 0.24 | -7.57 | 0.6 | 3.85 | 0.05 | -0.57 | 4.21 |
| Left | Humerus | Modern | BMOC | BMOC075 | -0.07 | -15.88 | 4.77 | 6.36 | 0.2 | -0.92 | 8.82 |
| Left | Humerus | Modern | BMOC | BMOC137 | 0.13 | -11.55 | -1.77 | 1.05 | -3.88 | -3.2 | 5.04 |
| Left | Humerus | Modern | BMOC | BMOC142 | 0.03 | -25.02 | -0.76 | 3.79 | 2.27 | -3.94 | 11.92 |
| Left | Humerus | Modern | BMOC | BMOC157 | | -14.85 | 2.84 | 8.33 | | -1.23 | 12.12 |
| Left | Humerus | Modern | BMOC | BMOC172 | -0.08 | 1.98 | -5.23 | 5.26 | -2.59 | -0.13 | 4.05 |
| Left | Humerus | Modern | BMOC | BMOC173 | -0.09 | -18.71 | -1.7 | 0.67 | -1.46 | -4.26 | 8.14 |
| Left | Humerus | Modern | BMOC | BMOC174 | 0.01 | -11.46 | -0.96 | 3.36 | -0.81 | -1.97 | 5.59 |
| Left | Humerus | Modern | BMOC | BMOC180 | 0.04 | 9.19 | 3.33 | 4.47 | -1.93 | 3.02 | 4.29 |
| Left | Humerus | Modern | BMOC | BMOC197 | 0.13 | -11.24 | 2.18 | 1.82 | -0.64 | -1.55 | 5.54 |
| Left | Humerus | Modern | BMOC | Fogel 2012 | -0.21 | -15.65 | 2.46 | 3.83 | -0.92 | -2.1 | 7.82 |

**Supplementary Table 6: Aggregate scores for all modern and subfossil Propithecus verreauxi individuals**

| *MEASURES* | *AGE* | *COLLECTION* | *SAMPLE ID* | *RIGHT FEMUR AVERAGE FOLD-CHANGE* | *LEFT FEMUR AVERAGE FOLD-CHANGE* | *RIGHT HUMERUS AVERAGE FOLD-CHANGE* | *LEFT HUMERUS AVERAGE FOLD-CHANGE* | *AGGREGATE SCORE* |
|---|---|---|---|---|---|---|---|---|
| 3D | Modern | BMOC | 550 | -0.041 | -0.07 | -0.051 | -0.079 | -0.06 |
| 3D | Modern | BMOC | 570 | -0.063 | -0.063 | -0.048 | -0.045 | -0.055 |
| 3D | Modern | BMOC | BMOC001 | -0.005 | 0.008 | -0.01 | 0.018 | 0.003 |
| 3D | Modern | BMOC | BMOC004 | -0.051 | -0.042 | -0.087 | -0.043 | -0.056 |
| 3D | Modern | BMOC | BMOC005 | 0.031 | 0.022 | NA | NA | 0.026 |
| 3D | Modern | BMOC | BMOC008 | 0.058 | 0.059 | 0.103 | 0.07 | 0.073 |
| 3D | Modern | BMOC | BMOC014 | 0.005 | -0.005 | NA | NA | 0 |
| 3D | Modern | BMOC | BMOC015 | 0.01 | -0.013 | 0.05 | 0.061 | 0.027 |
| 3D | Modern | BMOC | BMOC019 | 0.054 | 0.005 | 0.04 | 0.049 | 0.037 |
| 3D | Modern | BMOC | BMOC020 | -0.012 | 0.009 | -0.007 | -0.001 | -0.003 |
| 3D | Modern | BMOC | BMOC021 | -0.024 | -0.01 | NA | NA | -0.017 |
| 3D | Modern | BMOC | BMOC028 | 0.108 | 0.034 | -0.021 | -0.029 | 0.023 |
| 3D | Modern | BMOC | BMOC030 | 0.042 | NA | 0.02 | NA | 0.031 |
| 3D | Modern | BMOC | BMOC035 | 0.013 | NA | NA | NA | 0.013 |
| 3D | Modern | BMOC | BMOC043 | NA | NA | -0.051 | -0.008 | -0.03 |
| 3D | Modern | BMOC | BMOC073 | 0.004 | 0.011 | 0.042 | 0.016 | 0.018 |
| 3D | Modern | BMOC | BMOC075 | -0.04 | -0.031 | -0.012 | -0.029 | -0.028 |
| 3D | Modern | BMOC | BMOC125 | NA | NA | 0.058 | NA | 0.058 |
| 3D | Modern | BMOC | BMOC137 | 0.089 | 0.119 | 0.108 | 0.099 | 0.104 |
| 3D | Modern | BMOC | BMOC142 | 0.038 | 0.021 | 0.024 | 0.031 | 0.029 |
| 3D | Modern | BMOC | BMOC156 | -0.02 | NA | NA | NA | -0.02 |
| 3D | Modern | BMOC | BMOC157 | 0.004 | 0.024 | 0.009 | -0.019 | 0.004 |
| 3D | Modern | BMOC | BMOC163 | NA | 0.133 | NA | NA | 0.133 |
| 3D | Modern | BMOC | BMOC169 | 0.044 | NA | NA | NA | 0.044 |
| 3D | Modern | BMOC | BMOC172 | -0.039 | -0.055 | -0.026 | -0.023 | -0.036 |
| 3D | Modern | BMOC | BMOC173 | -0.03 | -0.018 | -0.037 | -0.016 | -0.025 |
| 3D | Modern | BMOC | BMOC174 | -0.013 | 0 | -0.008 | 0.007 | -0.004 |
| 3D | Modern | BMOC | BMOC180 | NA | 0.044 | 0.02 | 0.042 | 0.035 |
| 3D | Modern | BMOC | BMOC191 | NA | 0.026 | NA | NA | 0.026 |
| 3D | Modern | BMOC | BMOC197 | -0.093 | -0.047 | -0.052 | -0.033 | -0.056 |
| 3D | Modern | BMOC | Fogel 2012 | NA | NA | -0.004 | -0.042 | -0.023 |
| 3D | Subfossil | TAO | TAO-66-2 | NA | NA | 0.281 | NA | 0.281 |
| 3D | Subfossil | TAO | TAO-66-21 | 0.141 | NA | NA | NA | 0.141 |
| 3D | Subfossil | TAO | TAO-66-23 | NA | 0.093 | NA | NA | 0.093 |
| 3D | Subfossil | TAO | TAO-66-24 | NA | 0.032 | NA | NA | 0.032 |
| 3D | Subfossil | TAO | TAO-66-25 | 0.041 | NA | NA | NA | 0.041 |
| 3D | Subfossil | TAO | TAO-66-26 | 0.02 | NA | NA | NA | 0.02 |
| 3D | Subfossil | TAO | TAO-66-29 | 0.021 | NA | NA | NA | 0.021 |
| 3D | Subfossil | TAO | TAO-66-30 | NA | 0.105 | NA | NA | 0.105 |
| 3D | Subfossil | TAO | TAO-66-31 | NA | -0.173 | NA | NA | -0.173 |
| 3D | Subfossil | TAO | TAO-66-32 | 0.127 | NA | NA | NA | 0.127 |
| 3D | Subfossil | TAO | TAO-66-33 | 0.244 | NA | NA | NA | 0.244 |
| 3D | Subfossil | TAO | TAO-66-34 | NA | 0.139 | NA | NA | 0.139 |
| Caliper | Modern | BMOC | 550 | -0.034 | -0.04 | -0.03 | -0.041 | -0.036 |
| Caliper | Modern | BMOC | 570 | -0.013 | -0.011 | -0.047 | -0.05 | -0.03 |
| Caliper | Modern | BMOC | BMOC001 | -0.002 | -0.014 | -0.001 | -0.01 | -0.007 |
| Caliper | Modern | BMOC | BMOC004 | -0.018 | -0.017 | -0.031 | -0.023 | -0.022 |
| Caliper | Modern | BMOC | BMOC005 | 0.042 | 0.035 | NA | NA | 0.039 |
| Caliper | Modern | BMOC | BMOC008 | 0.054 | 0.044 | 0.072 | 0.038 | 0.052 |
| Caliper | Modern | BMOC | BMOC014 | -0.002 | -0.066 | NA | NA | -0.034 |
| Caliper | Modern | BMOC | BMOC015 | -0.022 | -0.025 | 0.027 | 0.02 | 0 |
| Caliper | Modern | BMOC | BMOC019 | 0.028 | 0.055 | 0.042 | 0.031 | 0.039 |
| Caliper | Modern | BMOC | BMOC020 | -0.02 | -0.016 | 0.003 | -0.016 | -0.012 |
| Caliper | Modern | BMOC | BMOC021 | 0.018 | -0.005 | NA | NA | 0.006 |
| Caliper | Modern | BMOC | BMOC028 | -0.003 | 0.021 | -0.02 | -0.033 | -0.008 |
| Caliper | Modern | BMOC | BMOC030 | 0.046 | NA | 0.041 | NA | 0.044 |
| Caliper | Modern | BMOC | BMOC035 | 0.001 | NA | NA | NA | 0.001 |
| Caliper | Modern | BMOC | BMOC043 | NA | NA | -0.002 | 0.005 | 0.002 |
| Caliper | Modern | BMOC | BMOC073 | 0.002 | -0.011 | -0.003 | 0.015 | 0.001 |
| Caliper | Modern | BMOC | BMOC075 | -0.018 | -0.023 | 0.032 | 0.394 | 0.096 |
| Caliper | Modern | BMOC | BMOC125 | NA | NA | 0.057 | NA | 0.057 |
| Caliper | Modern | BMOC | BMOC137 | 0.077 | 0.079 | 0.085 | 0.062 | 0.076 |
| Caliper | Modern | BMOC | BMOC142 | 0.039 | 0.023 | 0.025 | -0.002 | 0.021 |
| Caliper | Modern | BMOC | BMOC156 | 0.029 | NA | NA | NA | 0.029 |
| Caliper | Modern | BMOC | BMOC157 | 0.027 | 0.026 | -0.028 | -0.041 | -0.004 |
| Caliper | Modern | BMOC | BMOC163 | NA | 0.099 | NA | NA | 0.099 |
| Caliper | Modern | BMOC | BMOC169 | -0.098 | NA | NA | NA | -0.098 |
| Caliper | Modern | BMOC | BMOC172 | -0.05 | -0.047 | -0.064 | -0.018 | -0.045 |
| Caliper | Modern | BMOC | BMOC173 | -0.015 | -0.017 | -0.022 | -0.045 | -0.025 |
| Caliper | Modern | BMOC | BMOC174 | 0.007 | 0.011 | -0.007 | -0.014 | -0.001 |
| Caliper | Modern | BMOC | BMOC180 | NA | 0.018 | 0.004 | 0.03 | 0.018 |
| Caliper | Modern | BMOC | BMOC191 | NA | -0.006 | NA | NA | -0.006 |
| Caliper | Modern | BMOC | BMOC197 | -0.033 | -0.033 | -0.026 | -0.035 | -0.032 |
| Caliper | Modern | BMOC | Fogel 2012 | NA | NA | -0.047 | -0.06 | -0.054 |
| Caliper | Subfossil | TAO | TAO-66-2 | NA | NA | 0.192 | NA | 0.192 |
| Caliper | Subfossil | TAO | TAO-66-21 | 0.135 | NA | NA | NA | 0.135 |
| Caliper | Subfossil | TAO | TAO-66-23 | NA | 0.069 | NA | NA | 0.069 |
| Caliper | Subfossil | TAO | TAO-66-24 | NA | 0.059 | NA | NA | 0.059 |
| Caliper | Subfossil | TAO | TAO-66-25 | 0.09 | NA | NA | NA | 0.09 |
| Caliper | Subfossil | TAO | TAO-66-26 | NA | NA | NA | NA | NA |
| Caliper | Subfossil | TAO | TAO-66-29 | NA | NA | NA | NA | NA |
| Caliper | Subfossil | TAO | TAO-66-30 | NA | 0.057 | NA | NA | 0.057 |
| Caliper | Subfossil | TAO | TAO-66-31 | NA | NA | NA | NA | NA |
| Caliper | Subfossil | TAO | TAO-66-32 | 0.068 | NA | NA | NA | 0.068 |
| Caliper | Subfossil | TAO | TAO-66-33 | NA | NA | NA | NA | NA |
| Caliper | Subfossil | TAO | TAO-66-34 | NA | 0.045 | NA | NA | 0.045 |
